# Defining conservation units in a highly diverse species: A case on Arctic charr

**DOI:** 10.1101/2025.05.12.653098

**Authors:** Sam Fenton, Colin W. Bean, Samuel A.M. Martin, Samuel J. Poultney, Antony Smith, Elvira de Eyto, Kathryn R. Elmer, Colin E. Adams

## Abstract

Defining appropriate conservation units is crucial to the protection and management of biodiversity. These delineations deliver further benefit when they include assessments of population vulnerability to extinction from pressures such as climate change. However, delineations and vulnerability assessments are particularly difficult within highly diverse species, such as the salmonid fish Arctic charr (*Salvelinus alpinus*), that show extensive phenotypic and genetic variation within and across locations, variable and complex life histories, and broad geographic distributions. As yet, the nature and scope of Arctic charr diversity has not been characterised at the scale needed to delineate key conservation units in Scotland. To identify evolutionarily significant and vulnerable populations to prioritise for conservation, we conducted a genomic study of Arctic charr populations across Britain and Ireland with a focus on Scottish populations (N=64 populations; 24,878 SNPs; 410 individuals). We found that most lake populations represented distinct genetic clusters, with limited gene flow between them and resulting in substantial genetic differentiation. Higher level groupings of genetic similarity across catchments likely reflect historic anadromy and migration, with populations primarily grouping east or west of the central watershed divide in Scotland. Analysing genetic offset, also known as genomic vulnerability, we identified strong inverse correlations between genetic vulnerability with latitude and distance to the sea suggesting that more southern and more inland populations are more vulnerable to loss due to climate change. Additionally, patterns of vulnerability across several additional metrics identified further populations that may be at high vulnerability to loss. We further used our genetic data, along with phenotypic and geographic information, to identify populations of greatest evolutionary significance. This highlighted the most important ones to protect are those in locations with multiple ecotypes, a key facet of functional Arctic charr biodiversity, and populations that are the only ones in their Hydrometric Area.

## Introduction

The management and protection of biodiversity requires both the delineation of appropriate units of conservation and an assessment of their vulnerability to damage or loss. While conservation action is frequently directed at the species level, an increasing body of work has demonstrated the importance of intraspecific, within-species, diversity to ecological versatility, ecosystem recovery, and adaptation to changing environments (Des Roches et al., 2018; Le Coeur et al., 2021; Reusch et al., 2005; Schindler et al., 2010). For many species, particularly highly diverse ones, conservation management at a species level is insufficient for protecting vital variation and it is becoming increasingly common practice to define intraspecific conservation units (Forester et al., 2022; Mee et al., 2015). However, there are many different functional definitions of what constitutes a biologically meaningful conservation unit, and the criteria used to define these units can drastically impact the assessment of conservation importance (Mee et al., 2015; Minter et al., 2021; Mussmann et al., 2020). For example, the emphasis on morphological characteristics in the Kottelat & Freyhof (2007) review of the freshwater fishes of Europe led to a number of highly debated species classifications (Adams & Maitland, 2007; Barthelemy et al., 2023; Crotti et al., 2020; Denys, 2021; Etheridge et al., 2012; Gratton et al., 2014; Vavalidis et al., 2019). As such, an approach is needed to delineate appropriate units of conservation based on genetics rather than inconsistent morphological traits, particularly on the intraspecific level.

With the aim of conserving the extant diversity of a species and protecting its adaptive potential (i.e. future biodiversity), definitions of intraspecific conservation units focus on identifying reproductively isolated groups, or groups showing high genetic differentiation as a proxy for reproductive isolation (Allendorf et al., 2022; Mable, 2019). Two of the most commonly applied conservation delineations are Evolutionarily Significant Units (ESUs) and Management Units (MUs) (Allendorf et al., 2022; Funk et al., 2012; Moritz, 1994). Broadly an ESU can be defined as a reproductively isolated population, or group of populations, that show notable adaptive differentiation or functional diversity for the species. However, a number of variations of this definition exist, with some placing emphasis on contemporary patterns of diversity while others focus more on distinct historical and evolutionary lineages (Casacci et al., 2014; Coates et al., 2018; De Guia & Saitoh, 2007; Moritz, 1994; Waples, 1991). In contrast, MUs are generally defined as demographically independent, reproductively isolated units without the element of adaptive differentiation used for ESUs (Funk et al., 2012; Moritz, 1994). Many studies define these units hierarchically, with a species consisting of several larger ESUs that themselves contain multiple MUs (Funk et al., 2012; Shaney et al., 2020). However, this is not always the case, as a species may comprise a single ESU, an ESU may contain a single MU within it, for example when an ESU consist of a sole population, and MUs may be delineated first with the candidacy of ESU status at different hierarchies then evaluated (Casacci et al., 2014; Forester et al., 2022).

Several data types, for example genetic, phenotypic, and geographic data, should be combined where possible when defining units of conservation (Hoelzel, 2023). Information from different data types can provide important evidence that may reinforce the conclusion from another; this idea of concordance was a key part of the original definition of an ESU (Ryder, 1986). In addition, different types of data can provide alternative insights into what is evolutionarily significant, for example when high morphological differentiation is not matched by high genetic differentiation (Apolônio Silva de Oliveira et al., 2017; Funk et al., 2012). Nonetheless many studies defining intraspecific conservation units have used a criterion based solely on genetic data (Robertson et al., 2014). The advent of large-scale genomic datasets that use both neutral and adaptive loci have increased capacity to delineate conservation units more precisely and with greater biological depth (Allendorf et al., 2022; Funk et al., 2012). How conservation units are delineated are highly susceptible to whether patterns of differentiation are considered at a genome-wide scale or restricted to single or several adaptive genes (Waples et al., 2022; Waples & Lindley, 2018).

Beyond the delineation of conservation units, another important requirement of conservation management is the need to identify habitats and sites that may require urgent conservation action or monitoring because they are more vulnerable to loss (Campbell et al., 2017; Sgrò et al., 2011). This can involve investigating both the population for conservation and its supporting habitat. For example, small habitats with lower carrying capacity will limit the size of a population and potentially its overall diversity, restricting its ability to respond to change (Fischer & Lindenmayer, 2007).

Increasing temperatures are a threat to many species, with increased water temperature shown to affect the development and morphology of freshwater species (Campbell et al., 2021; Esin et al., 2021; Muir et al., 2022). The threat levels are magnified for populations in smaller lakes or streams which may have less refuge from high water temperatures when compared with populations in larger habitats (Campbell et al., 2017; Isaak & Young, 2023). Thus, using the combination of population genetics and environmental information to identify the level of the threat to a population from various factors can help identify not just vulnerable species but also vulnerable sites for that species to prioritise for conservation action.

A species that aptly demonstrates the complexity of defining units of conservation is the Arctic charr (*Salvelinus alpinus*), a salmonid fish. Across its Holarctic range, the species exhibits very high variation in the expression of phenotypic traits and high levels of genetic variation (Gordeeva et al., 2015; Jonsson & Jonsson, 2001; Klemetsen, 2010; Maitland & Adams, 2018; Recknagel et al., 2017a; Wilson et al., 2004). Across its distribution, there are a number of lakes that contain multiple distinct populations, known as ecotypes, diverged to specialise in different trophic niches and living in sympatry (Elmer, 2016; Klemetsen, 2010; Maitland & Adams, 2018). These ecotypes are often distinguished by differences in their diet and foraging tactics but can be differentiated in other characters such as morphology and spawning time (Adams et al., 1998; Doenz et al., 2019; Garduño-Paz et al., 2012; Hooker et al., 2016; Jacobs et al., 2020; Kess et al., 2021; Malmquist et al., 1992; Skúlason et al., 1996). The presence of distinct sympatric ecotypes tends to be associated with local environmental features such as lake size and climate (Blain et al., 2025; Fenton et al., 2024a; Tiddy et al., 2024). The extent of divergence between co-existing sympatric ecotypes varies by location and each represents an important example of the functional diversity of the species (Adams et al., 2007a; Adams & Huntingford, 2004; Fenton et al., 2024b; Gordeeva et al., 2015; Moccetti et al., 2019; Schluter, 1996). Many of these instances of ecotype divergence are the result of secondary contact and admixture of multiple lineages that colonised post-glaciation, while others represent recent sympatric divergences from a single lineage and therefore show lower levels of genetic differentiation but frequently clear differentiation in phenotypes (Garduño-Paz et al., 2012; Jacobs et al., 2020; Kettle-White, 2001).

The high intraspecific diversity of Arctic charr has resulted in both the species and their habitats being designated as having high conservation, natural heritage, and scientific value for many countries, including the UK (Bean et al., 2018). In Britain and Ireland, the species is not anadromous, remaining in freshwater throughout the life cycle (Finstad & Hein, 2012; Jørgensen & Johnsen, 2014; Maitland & Adams, 2018) thereby inhibiting contemporary gene flow between populations in unconnected river catchments. This effect is compounded by the existence of multiple lakes which harbour sympatric and independently evolved ecotypes with genetic, phenotypic and ecological distinctiveness (Fenton et al., 2024b; Hooker et al., 2016; Jacobs et al., 2020). Given the natural heritage value and the geographic isolation of populations, there is a particular need to identify evolutionary significant and/or vulnerable sites for Arctic charr for conservation purposes. Recent guidelines for selection of Sites of Special Scientific Interest (SSSIs) in Great Britain make a strong case for selection and protection of sites supporting within-species diversity and seeks to preserve the habitats giving rise to this diversity (Bean et al., 2018). There is also a concerted movement towards the protection of genetic diversity and future adaptive potential of species, particularly those known to threatened, both in the UK and internationally in response to Aichi Targets (2011-2020) and the Kunming-Montreal framework (2022-2030) (Department of Agriculture et al., 2025; Hoban et al., 2025; Hollingsworth et al., 2019). However, given that management resources are limited, it is important to define populations, or groups of populations, that are arguably higher priority for protection than others.

To delineate conservation units and identify vulnerable populations, we conducted a national scale genomic study of Arctic charr in Scotland, contextualised across Britain and Ireland. We evaluated multiple approaches to determine populations for conservation action using a high-density genome-wide dataset of SNPs. We analysed overall patterns of genome-wide genetic structuring and phylogenetic analysis across all populations and catchments. Further we examined patterns of variation in putatively adaptive SNPs associated with lake environment. We inferred populations that we predict may be of higher risk to loss in the future, via explicit vulnerability metrics such as genetic offset (genomic vulnerability). We then used our analyses in tandem with phenotypic and ecological knowledge, and geographical information, to delineate ESUs and MUs within the species and highlight populations of significance to protect.

## Materials and methods

### Sample collection

Arctic charr were sampled from 49 Scottish lakes, three Irish lakes, two Welsh lakes, and one English lake (Figure 1) between 1997 and 2021 using 30 m x 1.5 m Nordic multi-panel mesh survey gill nets (CEN, 2015) except at Llyn Padarn where fry traps were used to catch fry samples. Sample numbers and locations for each lake are indicated in Table 1. Eight of the sampled lakes in Scotland contain multiple sympatric ecotypes (Adams et al., 1998; Fraser et al., 1998; Jacobs et al., 2020). The dataset includes three instances of parapatric divergences (two neighbouring lakes from the same river system supporting two ecotypes): Shin-Merkland, Doine-Lubnaig, and Stack-More (Adams et al., 2006; Adams et al., 2007a). Each ecotype was considered its own population in all steps of data generation and analysis unless otherwise specified. Therefore, in this paper, we use the term population to refer to all Arctic charr individuals in the same lake of origin or a group of individuals that share morphological/ecological traits within a lake. Lochs Maree and Stack are both known to contain multiple ecotypes (Adams et al., 2008) however these were not sampled explicitly and so each of those lakes was considered as one population for the purposes of this study. In total, we had samples from 64 different Arctic charr populations covering 26 of the 27 Scottish Hydrometric Areas known to contain extant Arctic charr populations (Maitland & Adams, 2018). The dataset included 10 individuals per population for each unimodal population and five individuals per ecotype in multiple ecotype lakes, wherever possible (Table 1). Hydrometric Areas (HAs) are defined as single river catchments or several contiguous river catchments which are designated across the United Kingdom (National River Flow Archive, 2014).

**Figure 1:**
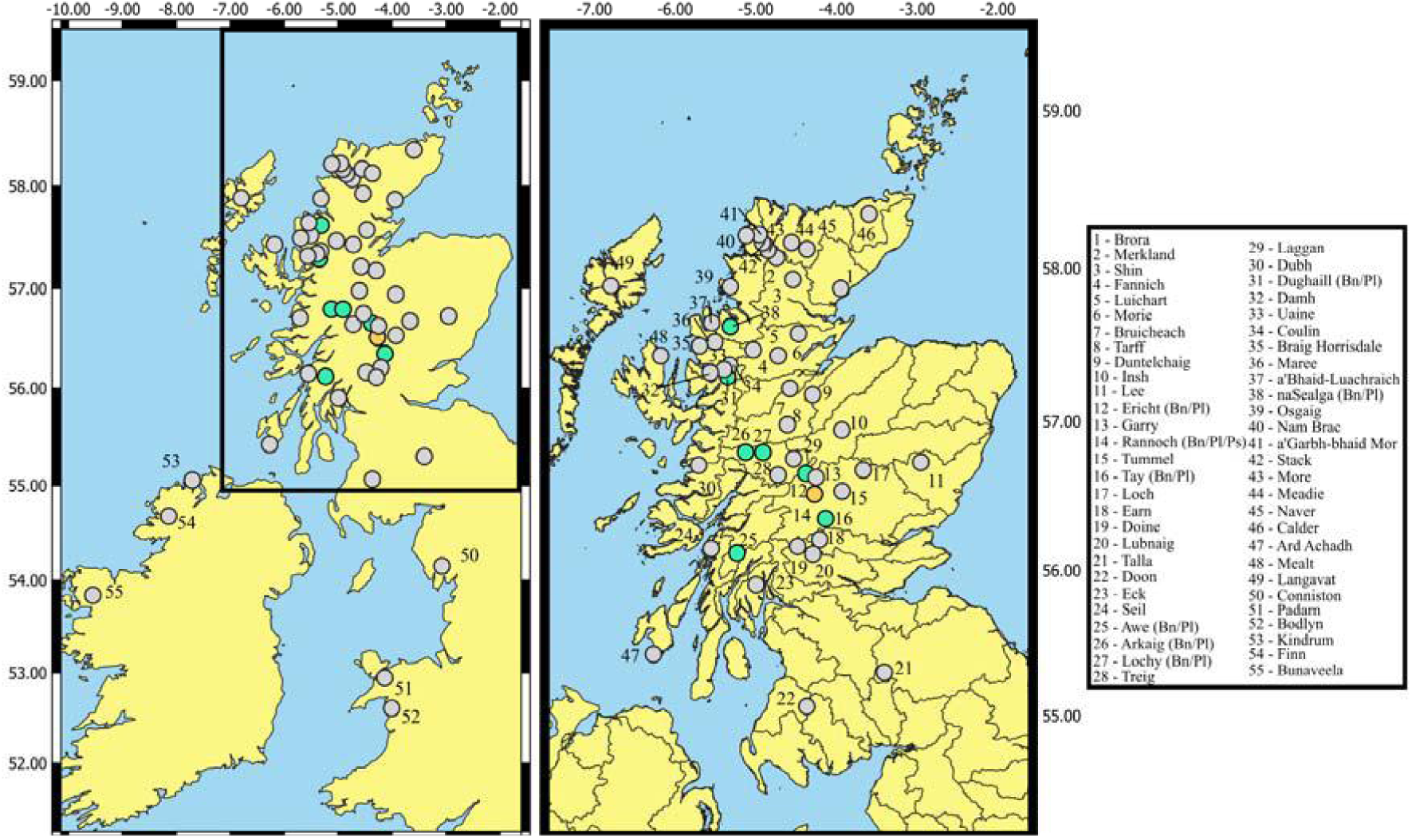
Map showing the location of the populations from the British Isles included in this study. Lake origin is indicated on the map in ascending number order following Hydrometric Area order. Lakes with multiple ecotypes are indicated by Bn/Pl/Ps delineators in the legend and with green (bimodal) and oranges (trimodal) colours respectively. Bn refers to benthivore, Pl refers to planktivore and Ps refers to piscivore. Hydrometric Area boundaries are indicated on the Scotland specific panel on the right.

**Table 1.** Overview of each of our 64 populations with summary statistics describing of genomic diversity. Location is either given as country (for outgroups) or Hydrometric Area for Scottish populations (numbered and named per the National River Flow Archive (2014). Pa refers to number of private alleles, He to expected heterozygosity, Ho to observed heterozygosity, π to nucleotide diversity, FST to population-specific FST.

| PopID | Latitude | Longitude | Sampling Year | N | Pa | Ho | He | $\pi$ | FST |
| --- | --- | --- | --- | --- | --- | --- | --- | --- | --- |
| a'Bhaid-Luachraich | 57.81 | -5.55 | 2012 | 10 | 32 | 0.093 | 0.075 | 0.080 | 0.512 |
| a'Garbh-bhaid Mor | 58.39 | -4.95 | 2021 | 10 | 80 | 0.065 | 0.048 | 0.051 | 0.688 |
| Ard Achadh | 55.36 | -6.16 | 2021 | 10 | 164 | 0.048 | 0.039 | 0.042 | 0.741 |
| Arkaig (Benthivore) | 56.97 | -5.14 | 2007 | 4 | 0 | 0.116 | 0.094 | 0.110 | 0.329 |
| Arkaig (Planktivore) | 56.97 | -5.14 | 2007 | 5 | 0 | 0.116 | 0.098 | 0.110 | 0.327 |
| Awe (Benthivore) | 56.30 | -5.23 | 1997 | 5 | 0 | 0.119 | 0.108 | 0.125 | 0.245 |
| Awe (Planktivore) | 56.30 | -5.23 | 1997 | 5 | 0 | 0.121 | 0.103 | 0.119 | 0.279 |
| Bodlyn | 52.80 | -4.01 | 2002 | 2 | 0 | 0.112 | 0.089 | 0.128 | 0.210 |
| Braig horrisdale | 57.67 | -5.67 | 2016 | 9 | 75 | 0.067 | 0.056 | 0.060 | 0.633 |
| Brora | 58.05 | -3.95 | 2010 | 4 | 1 | 0.095 | 0.074 | 0.088 | 0.461 |
| Bruicheach | 57.39 | -4.58 | 2009 | 8 | 24 | 0.075 | 0.059 | 0.065 | 0.601 |
| Bunaveela | 54.02 | -9.54 | 2001 | 10 | 127 | 0.054 | 0.044 | 0.047 | 0.714 |
| Calder | 58.52 | -3.59 | 2021 | 3 | 0 | 0.108 | 0.084 | 0.102 | 0.377 |
| Conniston | 54.34 | -3.07 | 2005 | 1 | 0 | 0.129 | 0.065 | 0.129 | 0.165 |
| Coulin | 57.55 | -5.35 | 1997 | 9 | 0 | 0.079 | 0.066 | 0.071 | 0.566 |
| Damh | 57.50 | -5.57 | 2008 | 2 | 0 | 0.112 | 0.069 | 0.104 | 0.355 |
| Doine | 56.34 | -4.48 | 1997 | 4 | 0 | 0.118 | 0.088 | 0.106 | 0.353 |
| Doon | 55.24 | -4.37 | 1997 | 6 | 0 | 0.112 | 0.090 | 0.100 | 0.387 |
| Dubh | 56.88 | -5.71 | 2021 | 5 | 12 | 0.124 | 0.091 | 0.108 | 0.339 |
| Dughaill (Benthivore) | 57.47 | -5.35 | 2013/2014 | 5 | 2 | 0.102 | 0.087 | 0.098 | 0.401 |
| Dughaill (Planktivore) | 57.47 | -5.35 | 2013/2014 | 5 | 0 | 0.105 | 0.093 | 0.105 | 0.361 |
| Duntelchaig | 57.35 | -4.29 | 2010 | 4 | 0 | 0.122 | 0.089 | 0.113 | 0.305 |
| Earn | 56.38 | -4.23 | 2016 | 9 | 27 | 0.102 | 0.083 | 0.089 | 0.453 |
| Eck | 56.08 | -4.99 | 1997 | 10 | 59 | 0.060 | 0.049 | 0.057 | 0.653 |
| Ericht (Benthivore) | 56.82 | -4.39 | 2007 | 4 | 1 | 0.127 | 0.106 | 0.126 | 0.229 |
| Ericht (Planktivore) | 56.82 | -4.39 | 2007 | 5 | 0 | 0.145 | 0.121 | 0.136 | 0.166 |
| Fannich | 57.64 | -4.96 | 2007 | 9 | 2 | 0.068 | 0.056 | 0.060 | 0.634 |
| Finn | 54.86 | -8.14 | 2012 | 1 | 0 | 0.051 | 0.025 | 0.051 | 0.679 |
| Garry | 56.80 | -4.25 | 2007 | 8 | 2 | 0.115 | 0.095 | 0.103 | 0.373 |
| Insh | 57.12 | -3.93 | 2007 | 10 | 8 | 0.092 | 0.077 | 0.082 | 0.498 |
| Kindrum | 55.23 | -7.71 | 1997 | 7 | 43 | 0.072 | 0.058 | 0.065 | 0.605 |
| Laggan | 56.92 | -4.55 | 2007 | 6 | 0 | 0.115 | 0.099 | 0.110 | 0.328 |
| Langavat | 58.04 | -6.82 | 1997 | 10 | 81 | 0.087 | 0.074 | 0.078 | 0.522 |
| Lee | 56.90 | -2.95 | 1997 | 9 | 35 | 0.108 | 0.090 | 0.096 | 0.413 |
| Loch | 56.85 | -3.67 | 2007 | 10 | 83 | 0.037 | 0.029 | 0.031 | 0.808 |
| Lochy (Benthivore) | 56.96 | -4.92 | 2009 | 2 | 0 | 0.188 | 0.124 | 0.171 | -<br>0.045 |
| Lochy (Planktivore) | 56.96 | -4.92 | 2009 | 5 | 0 | 0.123 | 0.104 | 0.116 | 0.288 |
| Lubnaig | 56.30 | -4.30 | 1997 | 10 | 2 | 0.103 | 0.078 | 0.100 | 0.387 |
| Luichart | 57.62 | -4.78 | 2009 | 10 | 29 | 0.102 | 0.086 | 0.092 | 0.440 |
| Maree | 57.71 | -5.53 | 2008 | 10 | 7 | 0.131 | 0.117 | 0.124 | 0.243 |
| Meadie | 58.33 | -4.56 | 2007 | 5 | 2 | 0.101 | 0.078 | 0.089 | 0.457 |
| Mealt | 57.61 | -6.18 | 2009 | 2 | 0 | 0.029 | 0.018 | 0.027 | 0.837 |
| Merkland | 58.23 | -4.73 | 1997 | 10 | 26 | 0.068 | 0.052 | 0.065 | 0.601 |
| More | 58.28 | -4.87 | 1998 | 4 | 0 | 0.121 | 0.091 | 0.113 | 0.309 |
| Morie | 57.75 | -4.47 | 2009 | 3 | 0 | 0.123 | 0.087 | 0.116 | 0.279 |
| Nam Brac | 58.38 | -5.12 | 2010 | 3 | 0 | 0.085 | 0.062 | 0.081 | 0.503 |
| naSealga (Benthivore) | 57.79 | -5.31 | 2015 | 5 | 0 | 0.089 | 0.071 | 0.083 | 0.497 |
| naSealga<br>(Planktivore) | 57.79 | -5.31 | 2015 | 5 | 0 | 0.088 | 0.072 | 0.084 | 0.492 |
| Naver | 58.30 | -4.35 | 2011 | 9 | 27 | 0.118 | 0.100 | 0.107 | 0.346 |
| Osgaig | 58.05 | -5.32 | 2015 | 9 | 35 | 0.086 | 0.073 | 0.079 | 0.517 |
| Padarn | 53.13 | -4.13 | 2020 | 10 | 65 | 0.095 | 0.082 | 0.087 | 0.469 |
| Rannoch (Benthivore) | 56.70 | -4.23 | 2010 | 1 | 0 | 0.101 | 0.050 | 0.101 | 0.382 |
| Rannoch (Planktivore) | 56.70 | -4.23 | 2010 | 6 | 5 | 0.115 | 0.098 | 0.109 | 0.215 |
| Rannoch (Piscivore) | 56.70 | -4.23 | 2010 | 4 | 0 | 0.133 | 0.111 | 0.128 | 0.336 |
| Seil | 56.32 | -5.55 | 2021 | 9 | 77 | 0.046 | 0.039 | 0.042 | 0.746 |
| Shin | 58.12 | -4.57 | 1997 | 7 | 1 | 0.111 | 0.095 | 0.104 | 0.364 |
| Stack | 58.34 | -4.92 | 1997 | 5 | 0 | 0.115 | 0.102 | 0.118 | 0.279 |
| Talla | 55.48 | -3.40 | 1997 | 8 | 0 | 0.098 | 0.084 | 0.091 | 0.444 |
| Tarff | 57.15 | -4.61 | 2012 | 8 | 57 | 0.049 | 0.038 | 0.042 | 0.744 |
| Tay (Benthivore) | 56.52 | -4.14 | 1997 | 5 | 1 | 0.127 | 0.105 | 0.122 | 0.257 |
| Tay (Planktivore) | 56.52 | -4.14 | 1997 | 5 | 1 | 0.121 | 0.107 | 0.123 | 0.254 |
| Treig | 56.81 | -4.73 | 2007 | 7 | 0 | 0.127 | 0.124 | 0.135 | 0.174 |
| Tummel | 56.71 | -3.95 | 2007 | 9 | 18 | 0.126 | 0.105 | 0.113 | 0.308 |
| Uaine | 57.52 | -5.39 | 1997 | 10 | 0 | 0.067 | 0.060 | 0.064 | 0.611 |

### DNA extraction and ddRADseq

DNA was extracted from fin clips, muscle, gill, head kidney, or from whole fry tissue using the Mackerey-Nagel NucleoSpin DNA Extraction Kit following manufacture recommendations. DNA quality and quantity were assessed using agarose gel electrophoresis and the Qubit Fluorometer with Life Technologies dsDNA BR assay. Four libraries of 90 samples plus five technical replicates were prepared using a ddRADseq protocol for Illumina, modified from Recknagel et al., (2015), and following Jacobs et al., (2020). Briefly, double restriction enzyme digestion was performed with *MspI* and *PstI*. A size selection range of 145-295bp was isolated using Pippin Prep. Final RAD-tag enrichment was performed over 10 PCR amplification cycles. Paired end 75bp sequencing was performed on the Illumina NextSeq500 platform at Glasgow Polyomics to generate 450 M reads per library (target of 4 M reads per individual and including 10% PhiX).

### Processing of ddRADseq data

Raw sequence quality was assessed using *FastQC v0.11.9* (https://www.bioinformatics.babraham.ac.uk/projects/fastqc/). The *process_radtags* pipeline in *Stacks v2.60* was used for demultiplexing raw sequence data (Rochette et al., 2019). To prevent a drop in sequence quality affecting mapping, the first 6 bp of the second read were removed using *Trimmomatic v0.39* (Bolger et al., 2014). New data were combined with previously generated data for eight Scottish lakes (Awe, Dughaill, na Sealga, Tay, Lubnaig, Merkland, Uaine, Eck) from Jacobs et al., (2020) (ddRADseq NCBI short read bioproject: PRJNA607173). Mapping was performed to the annotated *Salvelinus sp.* genome (ASM291031v2) using *bwa mem v0.7.16* using a seed length of 20 (Li, 2013). Reads with a mapping quality < 20 were filtered out using *samtools v1.15* (Danecek et al., 2021). Ten individuals with < 1M mapped reads were removed due to the high amounts of missing data. The *gstacks* module of *Stacks*, was run with the –gt-alpha set to 0.01 to ensure called genotypes were reliable when RAD-loci building. SNPs were retained in the populations module of *Stacks* if they met the following criteria: present in 66% of all individuals in each population and across all populations, a minimum minor allele frequency of 0.01, maximum observed heterozygosity of 0.5 with only the first SNP per locus retained. We used the script *filter_hwe_by_pop.pl* (https://github.com/jpuritz/dDocent) to filter any SNPs that were out of Hardy-Weinberg Equilibrium within populations, although no such SNPs were detected. *VCFtools v0.1.16* was used to identify and remove any SNPS with read depth <10X or >30x using the 2x average depth rule to determine allowed maximum depth (Crotti et al., 2021; Danecek et al., 2011) with our average depth being ∼15x.

### Genotyping error

We used an R script published by Mastretta-Yanes et al., (2015) to estimate the rate of genotyping error using 20 replicated individuals. Overall error rate was calculated to be 2.6%. The R script was then adapted to identify particular SNPs that showed a high rate of mismatching across technical replicates. SNPs that mismatched in more than 1 replicate pair (i.e. >5% genotyping error rate) were removed from the dataset for all individuals. Genotyping error was then recalculated from technical replicates and resulted in a mean error rate of 1.4%. Technical replicate samples were then removed from the dataset, retaining the replicate with the higher coverage. After filtering we had 24,878 SNPs and 410 individuals covering 64 different populations. While an upper limit of 34% missing data was allowed, average missing data per SNP was just over 10%, with roughly 4,600 SNPs showing 20% or more missing data.

### Summary statistics

Pairwise F_ST_, observed and expected heterozygosity, number of private alleles and F_IS_ were generated for all populations and loci using the –fstats parameter in the *populations* module of *Stacks*. Population specific FST was calculated using the *betas()* function in the *adegenet R package v2.1.10* (Jombart, 2008; Kitada et al., 2021).

### Identification of SNPs correlated to environmental and climatic variables

We conducted a Redundancy Analysis (RDA) using the *vegan v2.6.2* R package to identify SNPs associated with variation in environmental and climatic variables (Table S1); these represent our putatively adaptive SNP dataset to analyse patterns of adaptive differentiation (Capblancq et al., 2018; Oksanen et al., 2022). RDA was conducted at the individual level with individuals in the same lochs given the same environmental variables. Environmental data for 19 variables was collected from WorldClim using the raster R package *v3.5-15* using latitude and longitude co-ordinates (resolution 2.5 arcmin) (Fick & Hijmans, 2017; Hijmans & van Etten, 2012). Bathymetric data on lake surface area, mean and max depth, and altitude was collected for each lake from published work (Fenton et al., 2024a; Maitland & Adams, 2018; Murray & Pullar, 1910). An additional bathymetric variable, littoral zone percentage (defined as the area of the lake shallower than 5 metres depth) was calculated using available bathymetric maps and *ImageJ v1.50i* (Murray & Pullar, 1910; Schneider et al., 2012).

To create a dataset of uncorrelated variables, as collinearity can affect genotype-environment patterns (Ratner, 2009), a collinearity matrix was made in R for all environmental and bathymetric variables (Table S2). Variables with correlation coefficient (ρ) of < 0.7 were retained as independent variables and in cases where variables showed high correlation (ρ > 0.7) only one variable was retained (Layton et al., 2021) (Table S3). This resulted in a set of 10 uncorrelated variables (mean diurnal range, isothermality, maximum temperature of the warmest month, mean temperature of the wettest quarter, mean temperature of the driest quarter, annual precipitation, altitude, mean lake depth, lake surface area and the proportion of the lake comprising littoral zone (as %)). Missing data was replaced with the most common genotype at each SNP across all individuals to enable the RDA analysis (Kamvar et al., 2017).

### Genes and regions containing adaptive SNPs

To determine if our putative adaptive SNPs were related to functional processes, we investigated the genes and regions containing these SNPs. First, we compared the position of adaptive SNPs with a database of markers for quantitative trait loci (QTL) for various phenotypes from a range of salmonid species mapped to the *Salvelinus sp.* genome (Fenton et al., 2024b). Second, we compared SNP positions to all the annotated genes in the *Salvelinus sp.* genome (ASM291031v2) to identify genes containing associated SNPs. Both of these comparisons were carried out using *BEDtools v2.27.1* (Quinlan & Hall, 2010), with a limit of the SNP being within ± 100kbp for QTLs and ± 1kbp for genes to be considered. We then ran gene ontology term overrepresentation analysis (ORA) for any genes deemed to contain adaptive SNPs using the *TopGO v2.40.0* R package (Alexa & Rahnenfuhrer, 2020) using all genes containing any RAD loci from our ddRADseq data (associated or otherwise) as the comparison dataset. Results were summarised using REVIGO (Supek et al., 2011).

### Analysis of population structuring

For analysis of population structuring, we removed any non-neutral SNPs to more accurately reflect neutral evolutionary population genomic patterns (Funk et al., 2012). These were the putatively adaptive SNPs (1,104 loci) associated with environmental variables and SNPs in the top 5% of global F_ST_ scores, inferred from *hierfstat* (Goudet, 2005). This left a dataset of 22,763 SNPs. Admixture modelling was run using *PopCluster* for K=2 to K=64 with medium scaling and 10 replicates per K (Wang, 2022). The FSTIS and DLK2 estimators were used to infer the optimum number of K (Figure S1). Principal component analyses were run for all SNPs and associated SNPs using the *adegenet R package*, with any missing data replaced with the mean allele frequency for that SNP across all individuals.

Distance-based phylogenetic relationship between populations were constructed using an unrooted Neighbour-Joining (NJ) tree in the *poppr R package v2.9.3* (Kamvar et al., 2015). Any SNPs with missing data were removed to prevent an effect on the phylogenetic trees. This left a dataset of 5,949 SNPs for phylogenetic analyses. The tree was run with 1,000 bootstraps with no root. AMOVA was run using the *poppr* R package with a strata list indicating grouping in the N-J tree, Hydrometric Area of origin, and population and was run with 1,000 repeats.

### Calculating genetic offset

Future forecast climatic data was derived from the Coupled Model Intercomparison Project Phase 5 (CMIP5) under the Representative Concentration Pathways (RCP) scenarios RCP 4.5 and RCP 8.5. The RCP 4.5 scenario assumes CO_2_ emissions begin to decline by 2045 while RCP 8.5 assumes that CO_2_ levels will continue to rise throughout the 21^st^ century. SNPs associated with climatic variables were determined using an RDA as described earlier. We used climate associated SNPs to ensure genetic offset did not simply reflect genetic drift rather than adaptive signals (Láruson et al., 2022). To ensure gradientforest ran without errors, we removed any SNPs that showed low variability in allele frequency (5 or less different minor allele frequency values across all populations), leaving us with a dataset of 235 SNPs. Genetic offset was not calculated for Lough Finn and Coniston Water due to insufficient number (<2) of individuals. The analysis was also run only including populations having 5 or more individuals to determine the effect of small sample sizes on this analysis (Aguirre-Liguori et al., 2023). To model current genotype-environment relationships, *gradientforest R package v0.1.32* was used with 250 trees (Ellis et al., 2012). This model was then used to transform current and future climatic data based on their importance in explaining genomic variation with the Euclidean distance between these current and future values representing genetic offset. Due to the way *gradientforest* models genotype-environment relationships, a single genetic offset score was calculated for each lake meaning that there was a single score for lakes containing multiple ecotypes rather than one per ecotype.

### Lake sensitivity metric

Lake sensitivity scores for each lake in Scotland were taken from Maitland & Adams (2018). To summarise, sensitivity scores were calculated based on three bathymetric and location variables: Maximum depth of the lake, surface area of the lake, and lake altitude (Winfield et al., 2010). Deeper lakes were deemed to offer more refuge from high surface water temperatures, so were given a lower score. Similarly, higher altitude was deemed to provide some protection from thermal stress, so higher altitude was given a lower score. Small lakes will support smaller populations than larger ones and are impacted more by extreme climate events as a result (Fischer & Lindenmayer, 2007) so lakes with a smaller surface area were given a higher score. Scores for each variable were combined to give a score ranging from 3 to 11 with a score of 3 being a lake of least concern for population loss while 11 is of highest concern (Table S4).

## Results

### Summary statistics

Across the genomic dataset (N = 24,878 SNPs) from 410 individuals from 64 populations of Arctic charr, observed heterozygosity ranged from 0.029 to 0.188 (Figure S2). The Loch Lochy benthivore ecotype population was the most genetically diverse, with the highest observed heterozygosity (H_O_ = 0.188) while the Loch Mealt population was the least genetically diverse (H_O_ = 0.029) (Table 1). The number of private alleles ranged from 0 to 164. The Loch Ard Achadh (P_A_ = 164), Lough Bunaveela (P_A_ = 127), and Loch Loch populations (P_A_ = 83) had the highest numbers of private alleles. The number of private alleles negatively correlated with observed heterozygosity (P < 0.001, R^2^ = 0.329).

### Population structuring

A neighbour-joining tree broadly separated populations by drainage divide in Scotland (Figure 2). Nearly all populations in river systems that flow to the east coast of Scotland appear in one cluster, populations in west-flowing river systems in a second. The west flowing system cluster formed two major subgroups. All our outgroup populations from elsewhere in Britain and Ireland appear amongst the west-flowing system population cluster. Within these larger clusters, populations in the same or nearby river catchments/Hydrometric Areas were usually sister taxa but some sister taxa were also geographically distant populations such as lochs Eck and na Sealga.

**Figure 2:**
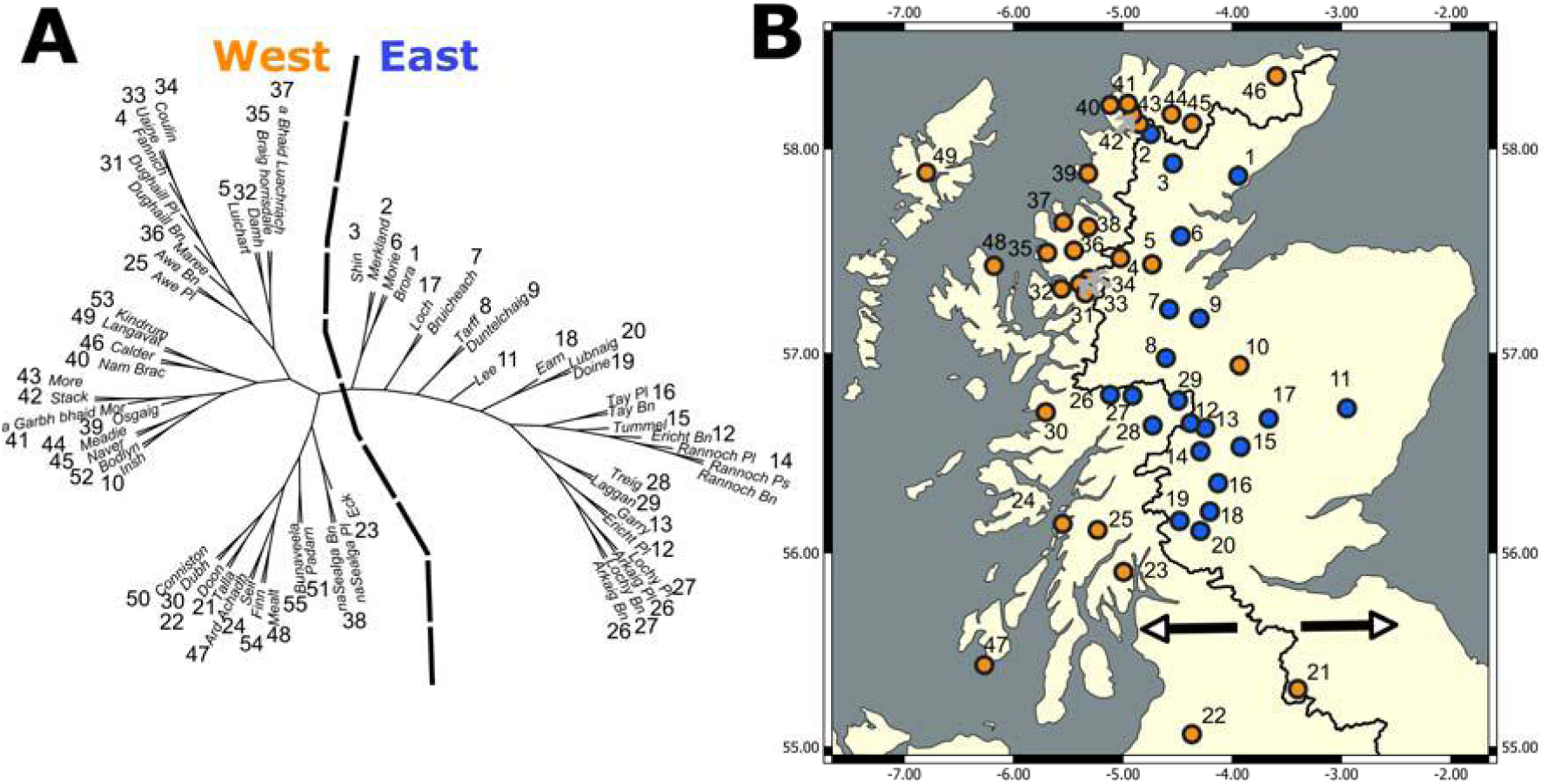
A) Neighbour-joining plot. NJ plot is split into largely east and west flowing river system groups and is indicated by the dotted line. B) shows the geographical distribution of this divide with the black line showing where the drainage divide occurs, populations to the left-hand side appear in west draining river catchments and those to the right in east draining. Populations are coloured by which side of the neighbour-joining tree they appear in, orange for west and blue for east. All outgroup populations are found in the west flowing group.

A principal component analysis did not separate major groups of populations, instead each PC axis only separated out one, or a small number of populations and explained a small amount of the total variance. Individuals from lochs Coulin, Uaine, Fannich, Maree, and the Dughaill planktivore populations in the northwest of mainland Scotland separate from all other populations along Eigenvector 1 (EV1) which explained 3.62% of the total variance (Figure 3A) while EV2 (explaining 2.57% of the total variance) splits the Loch Loch population, which is in central eastern Scotland, from all others suggesting high levels of population distinctiveness.

**Figure 3:**
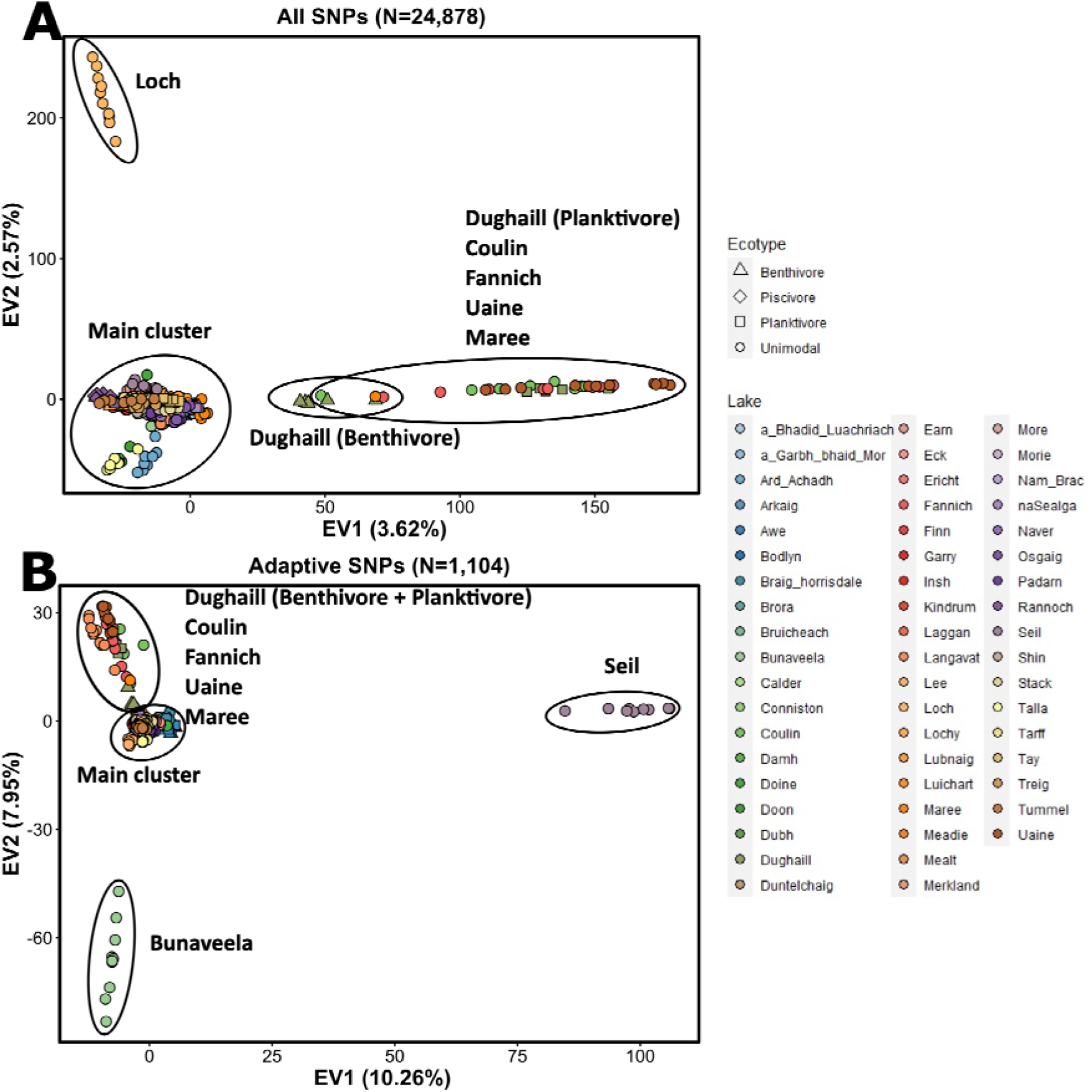
Principal components analyses based on all SNPs (A) (N=24,878) and associated SNPs (N=1,104) (B). Individual points are coloured by lake of origin indicated in the legend. Shape is based on population ecotype, triangle for benthivores, diamond for piscivores and squares for planktivores. Unimodal populations are given circles. EV refers to eigenvector.

Pairwise genetic differentiation was high between most populations, with a mean pairwise F_ST_ of 0.256 across all populations (Figure 4, Table S5). The lowest genetic differentiation F_ST_ (F_ST_= 0.045) was between Loch Doon and the Talla Reservoir population (which is a conservation refuge population derived from Loch Doon ancestors in the late 1980s (Maitland et al., 2007)). The highest pairwise differentiation was between Lough Finn in Ireland and the Loch Rannoch benthivore population (F_ST_ = 0.663), this is probably an artifact of the small number of individuals we had for each population. Several populations such as Loch Loch (mean pairwise F_ST_ = 0.384), Loch Seil (mean pairwise F_ST_ = 0.371), and Loch Tarff (mean pairwise F_ST_ = 0.357) showed particularly high pairwise F_ST_ in all comparisons (all pairwise F_ST_ > 0.2). Indeed, comparisons using population-specific F_ST_ showed these populations to be very high as well (F_ST_ > 0.7) (Table 1) (Kitada et al., 2021).

**Figure 4:**
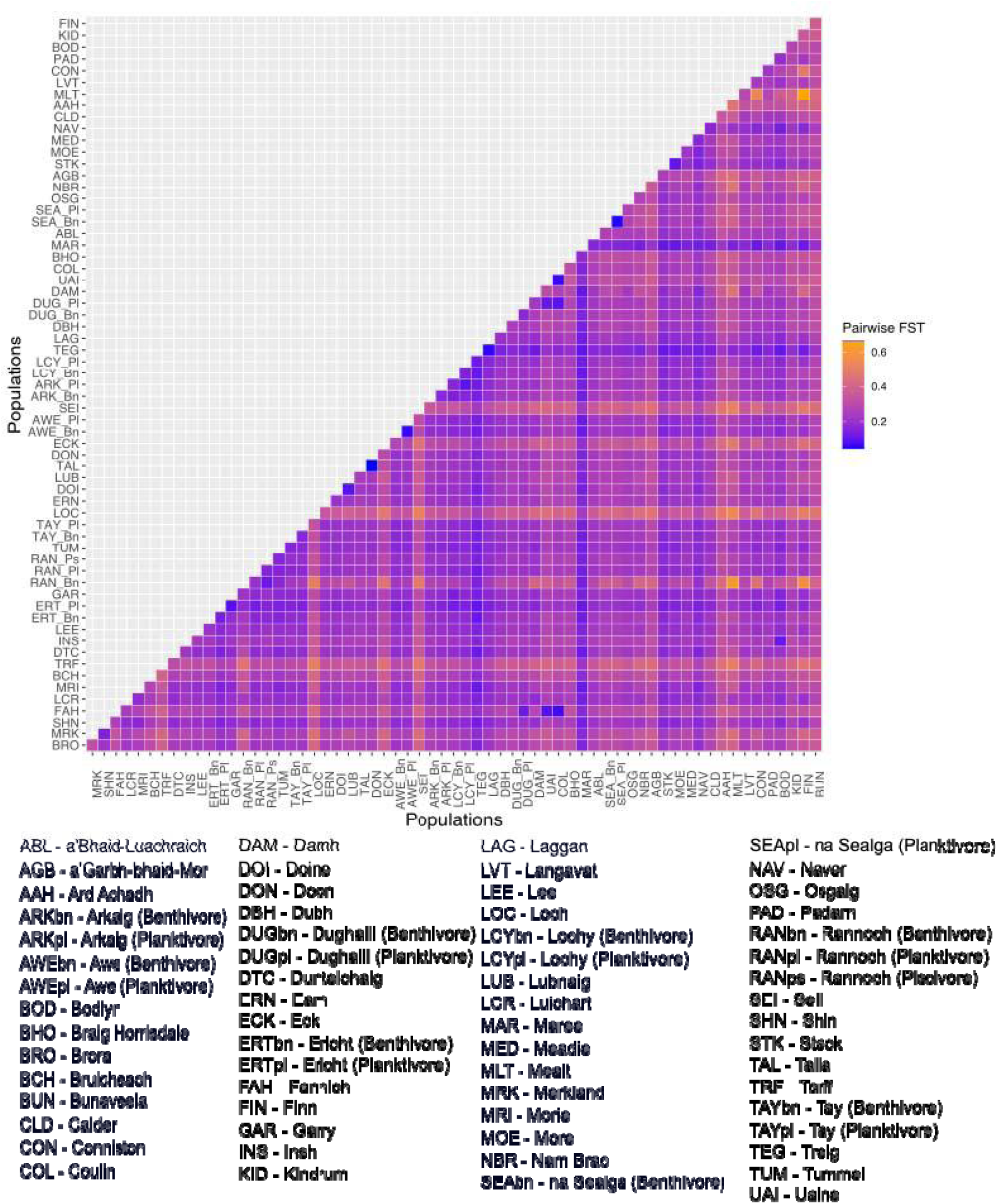
Plot of pairwise F_ST_ comparisons. F_ST_ scores are colours in a range of blue to orange going from low to high. Populations are order in ascending Hydrometric area of origin, order as indicated in Figure 1.

Admixture analysis was used to infer the number of genetic clusters (K) in the dataset. Fifty distinct clusters were identified as the most likely number in both DLK2 and FSTIS (Figure S1), with most of these clusters consisting of all individuals from a single population and evidence of admixture was rare (Figure 5). An analysis to partition the molecular variance found that most variation was explained by populations within Hydrometric Areas (35.4%). The east-west split found in the topology of the neighbour-joining tree explains <1% of the total variance in the AMOVA model. Hydrometric Area delineations within the east-west split explains 13.6% of the total variance. (Table S6).

**Figure 5:**
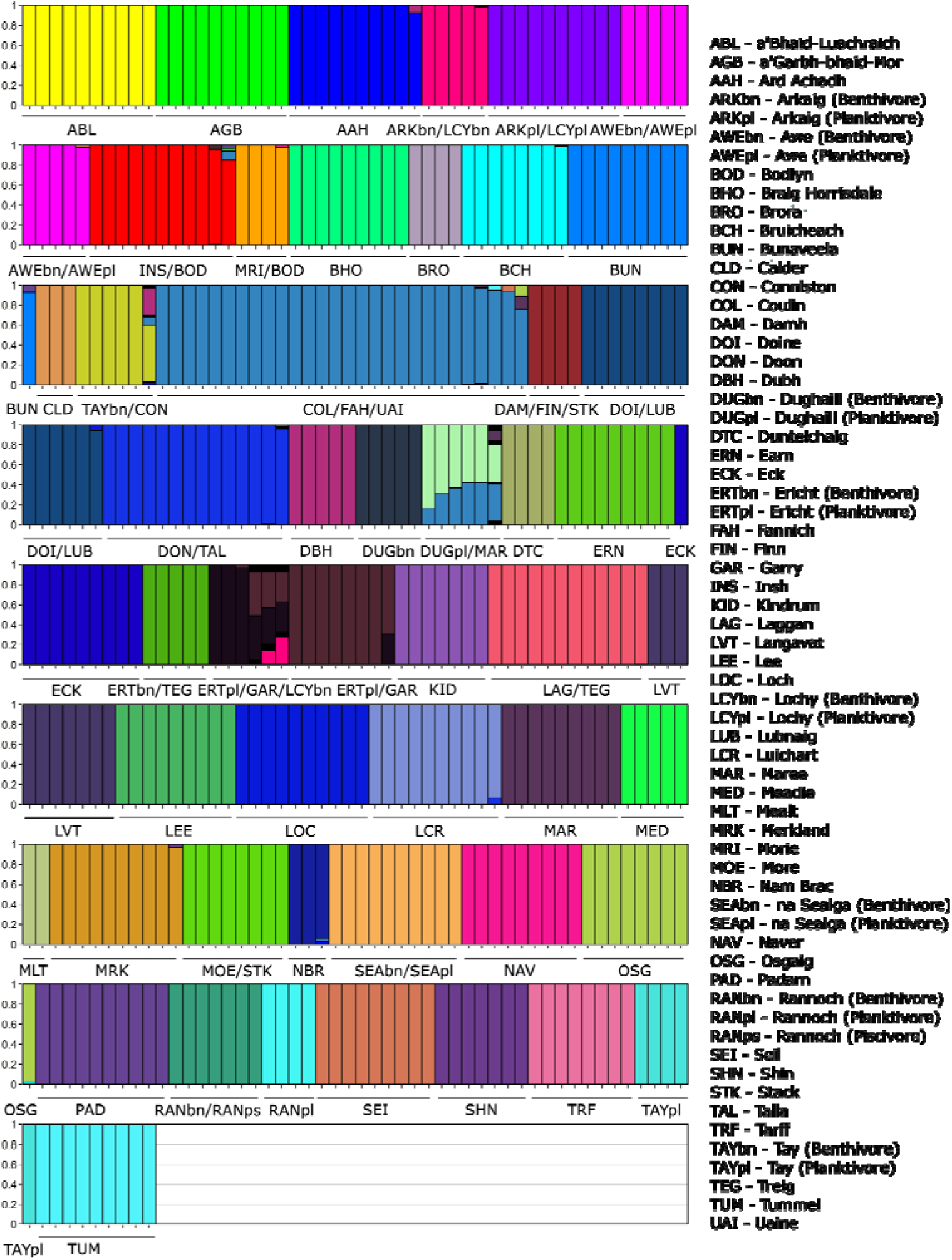
Admixture plot for most likely number of genetic clusters (K), K=50. Individuals are ordered by their genetic cluster. Most populations split into their own distinct genetic clusters with only some proximal populations sharing the same clusters. 3 letter codes for each population are provided. Ecotypes for relevant populations are indicated, bn for benthivore, pl for planktivore, ps for piscivore.

### Patterns of putatively adaptive variation

We used a redundancy analysis to identify SNPs associated with changes in climate, altitude, and lake bathymetry variables. Approximately 10% of the total variance in the whole dataset was explained by these environmental predictors. Bio3 (Isothermality (BIO2/BIO7)), Bio8 (Mean Temperature of Wettest Quarter), Bio9 (Mean Temperature of Driest Quarter) all counterposed in the opposite direction of Bio12 (Annual Precipitation) and Bio2 (mean diurnal range) on RDA1 (Figure S3). Altitude and littoral zone percentage counterposed in the opposite direction of mean lake depth and surface on RDA2.

Our analysis showed that all 10 constrained axes were significant in explaining variance and so we then identified SNPs with significant associations across all of them (Table S7). Overall, we identified a set of 1,104 SNPs as putatively adaptive to broad scale environmental conditions and lake bathymetry (Table S8). Surface area was the strongest predictor for the highest number of SNPs (N=234) followed by Bio9 (Mean Temperature of Driest Quarter) (N=166) (Table S9). This is reflected in the hierarchy of the importance of environmental variables in explaining adaptive variance with surface area the most important variable followed by Bio9 (Figure S4). When identifying the positions of the putatively adaptive SNPs to annotated genes in the *Salvelinus sp.* genome, we found that 586 known genes contained, or were proximal to, at least one of the 1104 putatively adaptive SNPs (Table S10). GO term analysis showed that amongst the known functions of these genes, a number of processes relating to formation, development, and morphogenesis such as skeletal system development (GO:0001501) and sagittal suture morphogenesis (GO:0060367) showed overrepresentation (Table S11).

When we compared positions to a database of the known genomic location of salmonid quantitative traits (Fenton et al., 2024b), we found that 35 QTLs contained, or were found close to, those loci we identified as adaptive SNPs (Table S12). Seven of these QTLs were originally identified from genetic mapping experiments from Arctic charr while 10 were from lake whitefish (*Coregonus clupeaformis*), 13 from coho salmon (*Oncorhynchus kisutch*), and five from lake trout (*Salvelinus namaycush*) all closely related species. Over half of the QTLs associated with morphological variation, with seven QTLs related to body length, six related to body shape, three related to body length, and a further three related to Fulton’s condition factor.

Exploring patterns of adaptive variation through a PCA revealed that EV1 explained 10.3% of the total variance and clearly differentiated the Loch Seil population from all other populations (Figure 3B). EV2 explained 8% of total variance and most clearly separates the Lough Bunaveela population (Ireland) from the other populations in one direction and a group including the lochan Uaine and lochs Coulin, Fannich, and Dughaill populations in the other direction. Conducting a PCA on the environmental variables, to determine if the patterns identified resulted from distinct environmental conditions, did not show the local conditions for these populations to be outliers in any obvious way (Figure S5).

### Susceptibility metrics

We assessed the potential vulnerability of populations of Arctic charr to a range of threats using three different metrics: genetic offset, lake sensitivity, and levels of observed heterozygosity focusing primarily on the Scottish populations.

We used genetic offset, also known as genomic vulnerability, as a measure of potential fitness maladaptation to changing climate, where higher values indicate elevated predicted maladaptation to future climatic change and risk of extirpation (Fitzpatrick & Keller, 2015). A redundancy analysis was used to identify a subset of SNPs (N=235) associated with change in climatic variables to be used to generate genetic offset scores. Under the RCP 4.5 emission scenario, genetic offset scores spanned from 0.005 (lowest risk) in the Loch Nam Brac population up to lochs Uaine (0.022) and Insh (0.028) (Figure 6). We did find a significant relationship between genetic offset scores and latitude, with southern populations generally showing higher genetic offset scores (P = 0.004, R^2^ = 0.13) (Figure 6A). However, many of the highest offset scores were found in the more northern parts of our distribution (Figure S6). We found a much stronger relationship between offset scores and distance to sea (P < 0.001, R^2^ = 0.23) with populations found in lakes at a further distance from the sea through river systems having higher genetic offset scores (Figure 6B). There was no significant relationship between genetic offset scores and observed heterozygosity (P = 0.249, R^2^ = 0.01) (Figure S7). Similar results were seen when using the more extreme RCP 8.5 scenario to calculate genetic offset scores with, for example, genetic offset scores showing a strongly correlated with latitude (Figure S8). The same patterns were seen when only calculating genetic offset with populations containing five or more individuals (Figure S9).

**Figure 6:**
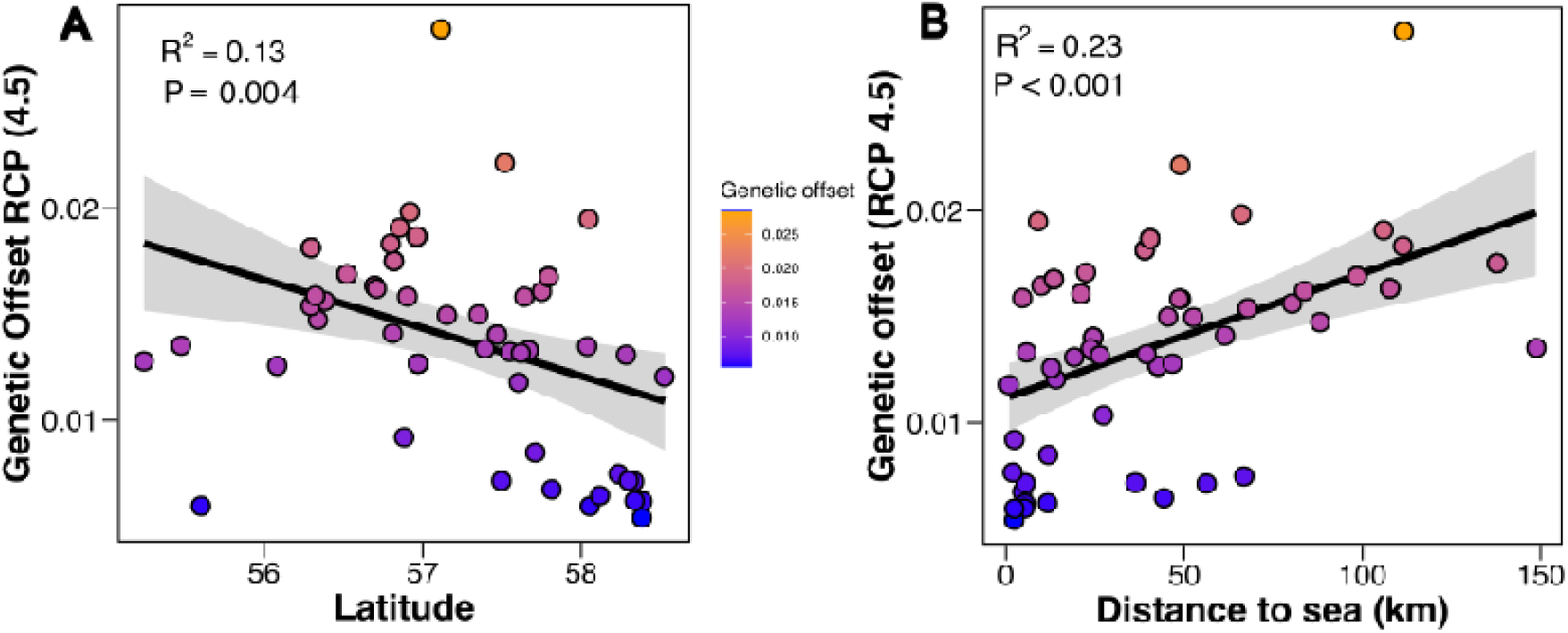
Genetic offset scores under RCP 4.5 scenario for each lake against A) latitude and B) distance from the sea in kilometres. Each point, population, is coloured by genetic offset score going from blue to orange as they go from low to high.

We assessed how the local environment affects potential population vulnerability using a metric of lake sensitivity based on lake bathymetry and altitude called lake sensitivity scores from Maitland and Adams, 2018. Possible scores ranged from 3 to 11, from least to most vulnerable. Three lochs, lochs Rannoch, Ericht, and Treig, had that lowest possible score of 3 in our dataset (Figure 7) whilst the highest score was 8, in lochs Seil and Ard Achadh, and the average score was 5. While the different metrics of susceptibility were largely uncorrelated with each other (Table S13), there were some populations such as Loch Seil that performed poorly across all metrics, showing low genetic diversity, a high lake sensitivity score, and a high genetic offset score (Figure 7). These populations were not found in any specific geographic region of our dataset and were spread out across the range.

**Figure 7.**
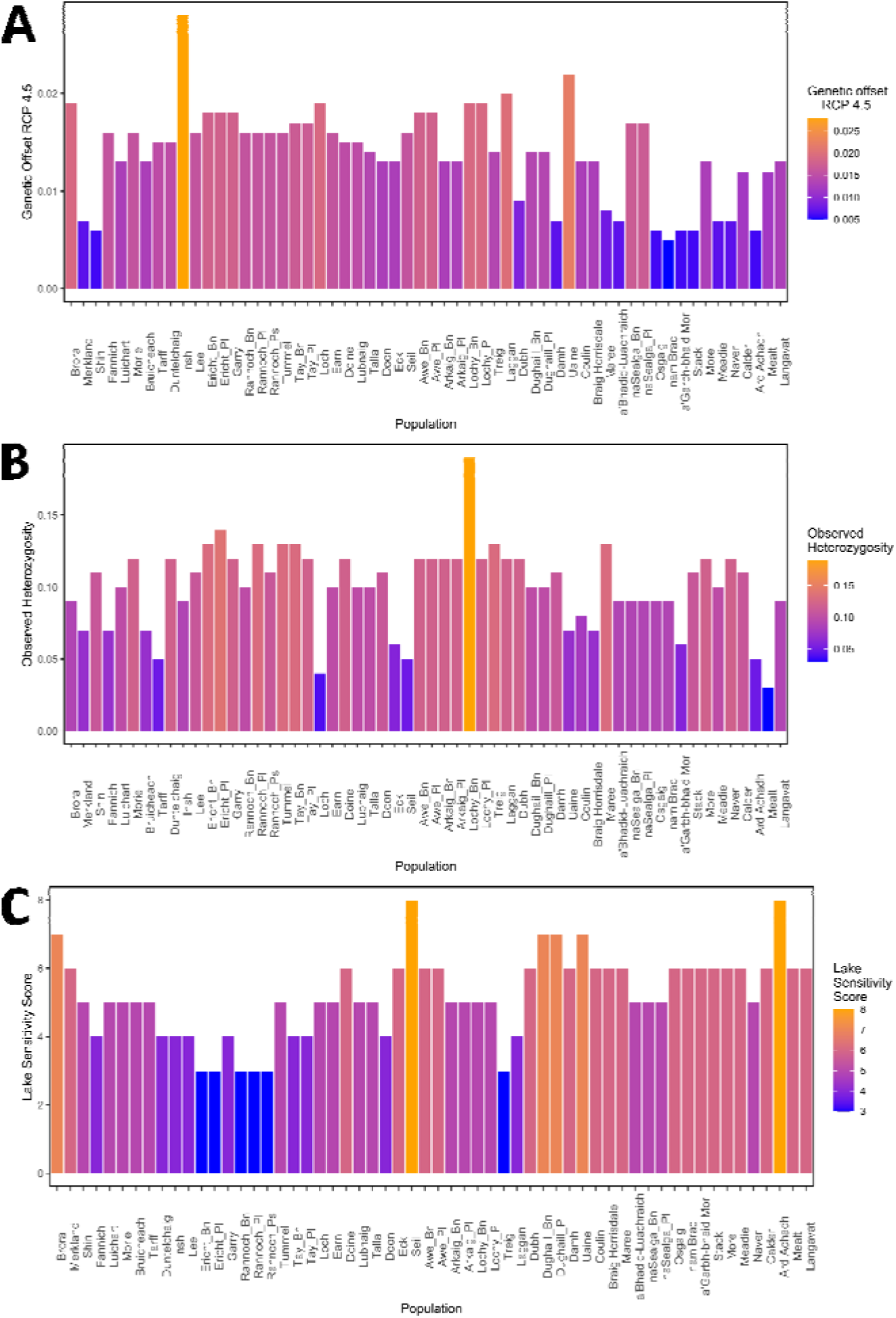
Plots showing the susceptibility score for genetic offset (A), heterozygosity (B) and lake sensitivity score (C) for each population in Scotland. Scores are coloured from blue to orange going from low to high values. Due to how each metric is generated, ecotypes in the same lake are given the same genetic offset and lake sensitivity scores. Observed heterozygosity was calculated based on all SNPs (N=24,878) while genetic offset was only calculated using SNPs associated with change in climatic variables (N=235). Lakes are listed along the x-axis in order of Hydrometric Area. Note that for genetic offset and Lake sensitivity scores, a higher score indicates higher vulnerability while for heterozygosity, a lower score indicates higher vulnerability.

## Discussion

Our results highlight how different genetic criteria to delineate priority populations for conservation can lead to differing classification of conservation units. We found that levels of genetic differentiation and genetic distinctiveness were high between almost all populations of Arctic charr in our study. The Hydrometric Area (the catchment or cluster of adjacent catchments) of origin of the population explains a small but meaningful component of the genetic variation which is otherwise very high across all populations. An important finding of our study further identified an important component of putatively adaptive variation that is unique to particular lakes and has the potential to result in differential responses between populations to future selection forces. Given the demonstrated high levels of population differentiation and weak geographic patterns, with distant geographic populations often showing similar levels of differentiation to geographical close ones, we believe geographically large-scale groupings would be inappropriate for conservation unit delineation. We instead focused on prioritising of lakes or regions that are conservation-relevant ESUs by exploring and balancing combined information and weighing different ecological and evolutionary components.

### National-scale structuring and regional genetic differentiation

At broader scales, an NJ tree largely clusters populations on either side of the hydrological central divide in Scotland forming east and west groups (Figure 2). While extant Arctic charr in Britain and Ireland are non-anadromous, the species was likely anadromous when it first colonised post-glaciation (Fenton et al., 2023). We expect that these east-west groupings likely represent a historic period of anadromy before the phenotype was lost with populations in nearby river systems clustering together due to the higher likelihood of contact with one another, a pattern known in freshwater fish species that do not disperse between unconnected river systems (Gómez & Lunt, 2007). These groupings influenced by drainage divide, are similar to that seen in North American populations of Brook trout (*Salvelinus fontinalis*) which can be found either side of the Eastern Continent Drainage Divide (Kazyak et al., 2022). However, this east-west grouping only explains a small proportion of the total variance (<1% as inferred from AMOVA) due to high levels of population differentiation and genetic distinctiveness (Table S6).

Analysis of population differentiation showed that most populations were quite differentiated. Many of the pairwise F_ST_ comparisons between populations were notably above the 0.15 value often used to determine significance of differentiation (Table S5) (Frankham et al., 2009). The patterns of high genetic differentiation are reflected in the admixture analysis with most populations forming their own distinct genetic clusters (Figure 5) and showing little support for clustering of these populations into larger groups. Populations showing lower genetic differentiation and some grouping at a level greater than populations were generally those found in interconnected lakes in the same catchment (e.g. lochs Doine and Lubnaig) or where populations are from the same lake (e.g. the ecotypes at lochs Awe or na Sealga). These findings are consistent with previous studies that found non-anadromous populations of Arctic charr generally show high levels of genetic differentiation and lower levels of genetic diversity (Salisbury et al., 2023; Shikano et al., 2015). In this Arctic charr are more structured and differentiated than many other salmonids such as Lake whitefish found in North America (Graham et al., 2022) and Brown trout (*Salmo trutta*) in both England and Ireland (Moccetti et al., 2024; Rodger et al., 2021).

A number of populations in our dataset are highly geographical isolated. Some of these lochs such as lochs Mealt and Ard Achadh, are on the Hebridean isles of Skye and Islay respectively and therefore isolated from mainland Scotland while other populations like Lough Bunaveela and Loch Seil are the only populations we have from their Hydrometric Area, and others like Loch Loch are highly separated from other populations in their Hydrometric area. We found that many of these isolated populations, such as Ard Ardchadh, Bunaveela, and Loch, tended to also have low heterozygosity with relatively high numbers of private alleles (Table 1). These isolated populations, that we expect have no contemporary gene flow with other populations, have gained unique mutations and their genetic differentiation may be driven more by genetic drift rather than selective pressures (Funk et al., 2016). The lack of gene flow reflected in the genetic distinctiveness of peripheral populations puts them at particular risk of extirpation, due to lower genetic diversity and higher risk of accumulation of deleterious mutations. Whether such isolated populations are of higher conservation value has been debated across species and depends on whether these populations exist in differing environments and their persistence potential (Finn et al., 2009; Lesica & Allendorf, 1995) and requires more investigation for these populations.

Several lochs in Scotland have sympatric and ecologically distinct ecotype pairs (Jacobs et al., 2020; Maitland & Adams, 2018). We found that levels of differentiation between ecotype pairs was variable and ranged from 0.05 to 0.20 pairwise F_ST_. The extent of differentiation is consistent with previously inferred differences in ecotypes evolutionary histories. Pairs that other research has concluded resulted from admixture after secondary contact of distinct colonising lineages showed higher differentiation (pairwise F_ST_ 0.16-0.20), such as the sympatric ecotype pairs in lochs Tay, Dughaill and the Rannoch planktivore versus the Rannoch piscivore/benthivore (Jacobs et al., 2020; Verspoor et al., 2010). In contrast the ecotypes that have been suggested to be sympatric divergences, particularly lochs Awe and na Sealga, have markedly lower genetic differentiation (pairwise F_ST_ 0.05-0.06) despite clear eco-morphological distinctiveness (Garduño-Paz et al., 2012; Jacobs et al., 2020). While less is known about the lochs Ericht, Lochy, and Arkaig ecotype pair origins because these divergences have not been studied in depth before, the levels of genetic differentiation we found here was relatively high (pairwise F_ST_ 0.14-0.17). We speculate that, given this level of differentiation, these pairs resulted from two post glacial invasions into the same lake and secondary contact, though this remains to be investigated further (Fenton, 2024)). Complex and differing evolutionary histories of sympatric ecotypes in postglacial lakes has been recently demonstrated in other areas, such as Arctic charr in Russia and European whitefish in Europe (Alekseyev et al., 2002; Fang et al., 2022; Öhlund et al., 2020) and may be a pervasive feature of contemporary populations even at nearby geographic scales.

### Adaptive differentiation

A potential route to delineating conservation units is to focus on adaptive genes (Hoelzel, 2023; Miller et al., 2024) as has been seen in Pacific salmonids in North America where conservation units are based on migration timing which is largely controlled by the *GREB1L/ROCK1* region and is an important phenotype to preserve variability in (Hess et al., 2016; Prince et al., 2017; Waples et al., 2022; Waples & Lindley, 2018). With this aim, we identified a set of 1,104 SNPs significantly correlated with climatic and lake variables across these populations to explore adaptive potential as well as investigate the genes and regions behind local adaptation across a broad scale. Given that previous studies on Arctic charr have highlighted that adaptive phenotypes, such as head and body shape in divergent ecotype populations, can be highly polygenic (Fenton et al., 2024b; Jacobs et al., 2020), we did not expect to define single regions that could be used conservation unit delineation. Indeed, we ended up finding a relatively high number of putatively adaptive SNPs and these are distributed across the genome (Table S8).

Over half of these environment-associated SNPs (N=624) were found within or proximal to known genes in the *Salvelinus sp.* genome. We found a high number of terms relating to morphogenesis and development as overrepresented amongst these SNPs. Fully half of the salmonid QTLs that we found contained putatively adaptive SNPs were related directly to morphology (Table S12). Further, 32 of the genes found to contain putatively adaptive SNPs were shared with a recent genome-wide association analysis that identified key genes that underlie head and body shape morphology differences between ecotypes of Arctic charr (Fenton et al., 2024b). While morphology was the most prevalent type of GO term in our dataset, several QTLs identified were related to other important phenotypes such as age of sexual maturation and spawning date, which are known to vary in our populations (Garduño-Paz et al., 2012). This suggests that a number of different aspects of phenotype may be key to local adaptation, although more data is required before concrete conclusion are drawn.

That we identified functional processes and genomic regions associated with head and bone morphology for SNPs associated with temperature variables is consistent with previous studies that indicate that both phenotypes are influenced by temperature across various fish species (Campbell et al., 2021; Hooker et al., 2023; Lema et al., 2019; Riesch et al., 2018). The QTLs containing environment-associated SNPs came from a number of different salmonid species, thereby indicating that key regions underlying important phenotypes are to a meaningful extent shared across species as has been previously suggested for salmonids (Fenton et al., 2024b; Jacobs et al., 2017; Salisbury & Ruzzante, 2022). Overall, adaptive variation is facilitated through a number of different processes and our results support the role of morphology acting as a key aspect of local adaptation across Arctic charr populations in the British Isles. The functional role of these candidate regions would be a valuable area of future research.

Patterns of differentiation in these putatively adaptive SNPs largely identified distinct single populations rather than larger groupings, particularly Loch Seil and Lough Bunaveela, which cluster separately (Figure 3B). A similar pattern was identified in similar study on Columbia spotted frogs, where a PCA only separate out single or small groups of populations form the larger distribution (Forester et al., 2022). The distinct populations at adaptive SNPs are not the same as the ones that appear as most distinct in the PCA using all SNPs (Figure 3A). The environmental conditions, lake bathymetry and climatic conditions for these populations were not obviously distinct from the other populations (Figure S5), and therefore the adaptive SNP divergence may be driven by parameters that were unquantified in our study or potentially due to genetic drift.

Our study is the first time many of these populations have been explored genetically and further evaluation is needed to properly understand the patterns seen. That we identified putatively adaptive SNPs located within genes and regions involved in ecologically relevant phenotypes is a notable finding and has implications for other species with highly isolated populations (Cortázar-Chinarro et al., 2017; Funk et al., 2016). However, caution is warranted on biological, environmental and analytical grounds. For example, it is possible that the putatively adaptive SNPs we identified involved in functional traits are not adaptive and could instead be maladaptive or non-beneficial in contemporary settings (Funk et al., 2012). Environmentally, WorldClim variables are atmospheric measures and it is not demonstrated how accurately they reflect the aquatic conditions of Arctic charr, given variation in lake depth, size and thermal buffering. Finally, high levels of neutral genetic structure can potentially confound gene-environment analysis and mimic signatures of local adaptation (Hay et al., 2022). As such we suggest further evidence is required before putatively adaptive SNPs are used to delineate important conservation decisions such as ESUs.

### Population susceptibility

A key part of conservation efforts is identifying populations that may be more vulnerable to loss and are therefore of higher conservation priority. Populations can be more vulnerable for a variety of reasons, from low levels of standing diversity to higher risk due to their environment, with these factors often uncorrelated with one another (Campbell et al., 2021; Fischer & Lindenmayer, 2007). Arctic charr is adapted to cold, highly oxygenated environments and as such the species is highly susceptible to climate change, particularly increasing water temperatures, and eutrophication (Kelly et al., 2020; Winfield et al., 2008). These risks are thought to be exacerbated at the southern edge of the species range, for example in North America and central Europe (Kelly et al., 2020; Layton et al., 2021). However, predicting the potential of population viability and risk is a challenging endeavour impacted by its environment and its population characteristics.

Genetic offset, often referred to as genomic vulnerability, has been suggested to be a metric for potential fitness maladaptation to future climatic change (Fitzpatrick et al., 2021; Layton et al., 2021). Several previous studies on genetic offset in salmonid species such as Arctic charr and Sockeye salmon *Oncorhynchus nerka* in North America have highlighted an important correlation between genetic offset scores and latitude with more southern populations tending to be at higher vulnerability and therefore needing to undergo greater magnitude of allelic change to survive in predicted future conditions (Layton et al., 2021; Tigano et al., 2024). Populations in Britain and Ireland are the southern extent of the European range, with the exception of remnant populations in the Alps (Tiberti & Splendiani, 2019). Therefore it is perhaps unsurprising that we found more southern populations may be more vulnerable to/require greater allelic change to survive future increase in temperature, which are predicted in the RCIP scenarios (Figure 6a) (Layton et al., 2021). The vulnerability of more southern populations is corroborated by the distribution across Britain, with only a few extant populations found in the southern parts of Scotland and northern England and Wales, and most of the recorded local extinctions occurring in these same regions (Maitland & Adams, 2018). However, the highest genetic offset scores we found were in the more northern latitudes, suggesting that genetic offset scores are not simply the product of latitude and the associated differences in climatic conditions and that other factors are contributing to genetic offset scores.

We found a much stronger relationship between genetic offset scores and geographic distance to the sea, with populations in lakes further to the sea having higher genetic offset scores (Figure 6B). This relationship suggests that more inland populations have had reduced levels of gene flow since colonisation, as seen in other fish species (Gouthier et al., 2023), and has resulted in them being more locally adapted but less flexible to future change. Notably, genetic offset scores were not driven by levels of observed heterozygosity, with heterozygosity used as an indirect measure of allele frequencies used in the gene-environment analysis, (Figure S7) suggesting scores were not driven by solely by standing levels of genetic variation (Láruson et al., 2022; Tigano et al., 2024).

It is worth noting that are several limitations of the genetic offset analysis (Láruson et al., 2022; Rellstab et al., 2021). Firstly, the small sample sizes for many populations means there is risk we did not accurately determine allele frequencies. Secondly, the inclusion of multiple ecotypes in the same lakes, and therefore have the same climatic variable values will introduce pseudo-replication and potentially bias the analyses. Additionally, as previously mentioned, WorldClim variable are atmospheric measures and so may not accurately reflect the lake environments.

While genetic offset and observed heterozygosity did not correlate in our dataset, observed heterozygosity still represents an important indicator of population vulnerability. Low levels of standing genetic variation suggests a population is less able to adapt to changes in their environment and are at higher risk of reduced population fitness due to inbreeding depression (Reed & Frankham, 2003). Similarly, populations in small lake environments may also be of concern for potential loss. Deeper, larger lakes hold more diverse populations both genetically and phenotypically (Fenton et al., 2024a; Recknagel et al., 2017), while small lakes have lower diversity and provide less refuge from the danger of rising water temperatures (Campbell et al., 2021; Fischer & Lindenmayer, 2007).

Overall, our metrics of susceptibility were largely independent of each other, allowing us to investigate vulnerability across multiple axes (Table S13) as is valuable in research focusing on determining conservation priority (Auber et al., 2022; Du et al., 2024). Looking across our metrics did not highlight any specific geographic regions of particular concern; the worst scoring populations in each metric were found distributed across our range (Figure 7). While we did note some populations that performed poorly across all metrics, such as Loch Seil, our analyses suggest that populations need to be assessed individually to determine their viability and vulnerability and there are no broader patterns such as an effect of Hydrometric Area. Our analyses do show however that the latitudinal position of a population, its distance from the sea, and overall lake size can also be used to give indications of potential vulnerability and can be used in a wider-scale analysis to identify potential regions of higher vulnerability or viability in the species distribution. Further work should look to include other potential threats to the species, such as lake acidification, hydroelectric work or non-native introductions to better assess population vulnerability (Knudsen et al., 2016; Maitland et al., 2007). Rather than trying to define a single definitive statistic to determine vulnerability, the investigation of vulnerability across a range of metrics can help us identify vulnerable populations in ways that may be missed by the use of a single metric.

### Delineation of ESUs and MUs

Conservation units like ESUs have had numerous definitions with the definition used often reflective of the system the unit is being applied to (Coates et al., 2018; De Guia & Saitoh, 2007; Moritz, 1994). Many studies that define ESUs and MUs delineate them in a hierarchical fashion, with larger ESUs containing multiple reproductively isolated populations considered to be MUs (Funk et al., 2012; Galland et al., 2021; Shaney et al., 2020). However, we argue that this dataset generated for the highly diverse Arctic charr is good example of how criteria should be best applied for the species and system in question. Given the patterns of our data, we suggest a non-hierarchical approach is more appropriate here. Arctic charr is listed as one of the priority fish species under the UK Biodiversity Framework (BRIG, 2007), in recognition of the need for conservation action to protect the species and the substantial diversity seen within it. Currently the conservation policy body for Scotland is focusing both on preservation of habitats-such as through Sites of Special Scientific Interest (SSSI), Special Area of Conservation (SAC), and National Nature Reserve (NNR) sites - and on genetic conservation units (GCUs) (Cavers et al., 2022). This study aims use evidence provided from conservation genomic analyses described herein to assess Arctic charr populations (including sympatric ecotype populations) for their relative conservation value in Scotland. Our findings feed into both the genetic conservation unit and the place-based perspective, given we examined both the genetic diversity and the role of local environment.

Given the high levels of genetic differentiation in our dataset, we created a logical framework for defining reasonable population groupings that we delineated into MUs and ESUs. We consider an ESU to be a population or group of populations that showed reproductive isolation to other groups and show evidence of adaptive differentiation, functional diversity and/or geographic importance (De Guia & Saitoh, 2007; Funk et al., 2012). We considered an MU to be populations or groups of populations that show reproductive isolation to one another (Funk et al., 2012; Moritz, 1994). This system is aligned with the rationale used for delineation of partial and full ESUs suggested by De Guia & Saitoh (2007) where partial ESUs show reproductive isolation but lack evidence of adaptive differentiation; although due to limited take up of the ‘partial ESU’ terminology, we proceed with the more standard use MUs in this case which have the same definition.

We do not rank MUs hierarchically within broader ESUs; the high levels of population differentiation seen in our admixture, F_ST_, and AMOVA analyses (Figures 4,5 and Table S6) limit the usefulness of delineating broader ESUs, which could be the major branches in our neighbour-joining tree for example (Figure 2). We also did not solely focus conservation priority on highly isolated populations with high numbers of private alleles, as we lack evidence that these are indeed adaptive differences or that these low diversity populations have high persistence potential (Finn et al., 2009; Lesica & Allendorf, 1995). We suggest that that populations in close proximity (<20km) and showing notable gene flow, conservation translocations (Maitland et al., 2007), and populations with notable shared genetic origin, e.g. recently diverged ecotypes (Jacobs et al., 2020), should be considered within the same ESU or MU. We consider the long-term conservation value for natural heritage to be higher of ESUs than MUs, as ESUs holds more adaptive and evolutionary potential (Funk et al., 2012; Moritz, 1994).

To delineate MUs, we first used the patterns of genetic differentiation, as a proxy for reproductive isolation, to determine the number of reproductively isolated populations or groups of populations (Allendorf et al., 2022; Palsbøll et al., 2007). Amongst the determined MUs, we then explore for evidence of substantial adaptive differentiation, functional diversity and/or geographic importance that warrants elevation to ESU status.

To delineate ESUs, we considered multiple lines of evidence, including phenotypic, environmental and geographic information, in addition to genetic data. Similar to our approach in exploring population susceptibility, where we choose not to focus on just one source of information, we merge as much evidence as possible (Waples et al., 2022; Waples & Lindley, 2018). We first explored patterns of adaptive differentiation through the identification of non-neutral and environment-associated SNPs, as other studies have done (Forester et al., 2022; Matala et al., 2014; Miller et al., 2024). While the identification of adaptive variance undeniably has importance in the delineation of conservation units (Hoelzel, 2023; Waples et al., 2022; Waples & Lindley, 2018), however as discussed earlier, we caution the use of the SNPs we identified as putatively adaptive for the basis of our conservation unit delineations. As such we then focused more on the existing phenotypic, environmental, and geographic information for these populations.

An aim of the current analyses is to provide an evidence base for the definition of high value polymorphic populations in Scotland. We deemed ecotype populations, either in multimodal lakes or parapatric divergences, as having high conservation value and should be considered ESUs due to their importance to the functional diversity of the species and its evolutionary future (Jonsson & Jonsson, 2001; Maitland & Adams, 2018) (Table 2). These includes ecotypes that do not show clear genetic differentiation but have reproductive isolation and ecological distinctiveness. These represent important incipient diverging groups, although not necessarily the same evolutionary potential as those resulting from secondary contact (Adams et al., 1998; Adams & Huntingford, 2004; Alexander & Adams, 2000; Garduño-Paz et al., 2010; Jacobs, 2018; Jonsson & Jonsson, 2001; Kettle-White, 2001). For example, Arctic charr in freshwater lakes known have very marked homing to their natal spawning sites, as seen in Lake Windermere (Frost, 1965), which potentially precipitates future divergences (Elmer, 2016). While some morphological and behavioural variation might be due to phenotypic plasticity (Hooker et al., 2023; Kristjánsson et al., 2018), the precautionary principle motivates us to nonetheless give some conservation priority to those populations. Populations that show notably lower genetic differentiation to an ecotype population but are not part of the system, i.e. not in the same lake or parapatric system, were placed in the same ESU. Populations that are the only extant population in their respective Hydrometric Area have a higher potential to gain unique and important adaptive variation, due to their extreme isolation, and their loss would significantly affect the geographic range of the species (Lesica & Allendorf, 1995) so we also considered these as ESUs.

**Table 2:** Conservation unit classifications for populations in our dataset as Evolutionarily Significant Units (ESUs) or management units (MUs). Delineation criteria for groups as ESUs are listed, in our study this was either due to be a diverging ecotype population or being ht eonly extant population in a Hydrometric Area.

| Population(s) | Conservation unit type | Delineation criteria for ESU |
| --- | --- | --- |
| a'Bhaid-Luachraich | MU |  |
| a'Garbh-bhaid Mor | MU |  |
| Ard Achadh | MU |  |
| Arkaig (Benthivore) | ESU | Diverged ecotype pop. |
| Arkaig (Planktivore) | ESU | Diverged ecotype pop. |
| Awe (Benthivore) | ESU | Diverged ecotype pop. |
| Awe (Planktivore) | ESU | Diverged ecotype pop. |
| Braig horrisdale | MU |  |
| Brora | MU |  |
| Bruicheach | ESU | Only extant HA pop. |
| Calder | MU |  |
| Coulin, Dughaill (Planktivore), Fannich, Uaine | ESU | Diverged ecotype pop. |
| Damh | MU |  |
| Doine | ESU | Diverged ecotype pop. |
| Doon, Talla | ESU | Only extant HA pop. |
| Dubh | MU |  |
| Dughaill (Benthivore) | ESU | Diverged ecotype pop. |
| Duntelchaig | MU |  |
| Earn | ESU | Only extant HA pop. |
| Eck | ESU | Only extant HA pop. |
| Ericht (Benthivore) | ESU | Diverged ecotype pop. |
| Ericht (Planktivore), Garry | ESU | Diverged ecotype pop. |
| Insh | MU |  |
| Laggan, Treig | MU |  |
| Langavat | MU |  |
| Lee | ESU | Only extant HA pop. |
| Loch | MU |  |
| Lochy (Benthivore) | ESU | Diverged ecotype pop. |
| Lochy (Planktivore) | ESU | Diverged ecotype pop. |
| Lubnaig | ESU |  |
| Luichart | MU |  |
| Maree | ESU | Diverged ecotype pop. |
| Meadie | MU |  |
| Mealt | MU |  |
| Merkland | ESU | Diverged ecotype pop. |
| More | ESU | Diverged ecotype pop. |
| Morie | MU |  |
| Nam Brac | MU |  |
| naSealga (Benthivore) | ESU | Diverged ecotype pop. |
| naSealga (Planktivore) | ESU | Diverged ecotype pop. |
| Naver | MU |  |
| Osgaig | MU |  |
| Rannoch (Benthivore) | ESU | Diverged ecotype pop. |
| Rannoch (Piscivore) | ESU | Diverged ecotype pop. |
| Rannoch (Planktivore) | ESU | Diverged ecotype pop. |
| Seil | MU |  |
| Shin | ESU | Diverged ecotype pop. |
| Stack | ESU | Diverged ecotype pop. |
| Tarff | MU |  |
| Tay (Benthivore) | ESU | Diverged ecotype pop. |
| Tay (Planktivore) | ESU | Diverged ecotype pop. |
| Tummel | MU |  |

The ecotype-polymorphic populations we consider to be ESUs are Awe (benthivore and planktivore), Arkaig (benthivore and planktivore), Doine, Dughaill (benthivore and planktivore), Ericht (benthivore and planktivore), Lochy (benthivore and planktivore), Lubnaig, Maree, Merkland, More, na Sealga (benthivore and planktivore), Shin, Stack, Rannoch (benthivore, planktivore, and piscivore), and Tay (benthivore and planktivore) (Table 2). The loch Dughaill planktivore ESU also contains the lochs Coulin and Fannich and lochan Uaine populations, due to patterns of genetic similarity seen in admixture and PCA analyses influenced by a shared ancestral history (Fenton, 2024). The known ecotypes at lochs Maree and Stack (Adams et al., 2008) should still be considered separate ESUs; though they were not directly studied in this work, they should be investigated in future.

ESUs for Arctic charr lakes that are the only populations in their respective Hydrometric Areas are in lochs Eck, Doon, Bruicheach, Lee, and Earn. We suggest the Loch of Girlsta, which is the sole extant population on the Shetland Islands, represents an ESU on the same criteria but as it was not included in our dataset it needs further investigation (Maitland & Adams, 2018).

We define all remaining groups with the working designations of MUs. In the scope of our study, this includes 24 populations that lie across the extent of Scotland (Fig 1, Table 2).

These designations we propose are well informed by genomic data and contemporary analyses, but should be considered recommendations about priority and not considered as definitive. More data on these populations should be used to further our understanding of the contemporary variation and relationship between populations, both within our dataset and those not currently covered, to help delineate conservation units. Further, there are many other charr populations in Scotland that are not in our dataset and would benefit from genetic characterising; Maitland & Adams (2018) lists 187 lochs with extant or likely extant populations of Arctic charr in Scotland. While the vulnerability of populations was not the focus of our conservation unit delineations, it is an important consideration for how to manage populations (Miller et al., 2024). Other factors such as unique historical lineages (Jacobsen et al., 2022; Moore et al., 2015) and adaptively important phenotypes, for example a difference in spawning (Walker, 2007), foraging/diet (Garduño-Paz & Adams, 2010) or habitat usage (Kristjánsson et al., 2012), would also be considered evolutionarily significant and worth prioritising for protection as ESUs (Zhao et al., 2020). Given that we now have an understanding of national-scale patterns of genetic differentiation and structuring, and thus we suggest how conservation units can be delineated, more focus should go into determining conservation priority from a management and policy perspective.

The focus on protecting environments and landscapes (Mainstone et al., 2018) already gives some protection to some Arctic charr populations in Scotland, irrespective of evolutionary genetics. For example, lochs Eck and Doon are included as notified features within protected sites (such as Sites of Special Scientific Interest) and are therefore protected in law (Bean et al., 2018; Mainstone et al., 2018). Lochs Doon, Lee, and Girlsta have also been flagged previously as sites that may be of higher risk of loss, due to factors such as lake acidification (Maitland et al., 1991, 2007). A number of lochs, such as Maree and Rannoch, are located within sites classified as Special Areas of Conservation under the EU Habitats Directive and Special Protection Areas (under the EU Birds Directive) to protect other habitats and species present, which also benefits Arctic charr (McLeod et al., 2009). However, it is evident that current landscape-based protection is not encompassing the ESUs and MUs defined here. Given the high diversity seen in Arctic charr, we urge that many of these ESU sites we have defined be considered for special protection, for example as SSSIs. While the formal designation of ESU population as a notified feature based purely on our findings is not always likely, inclusion of some ESUs populations to the list of notified features in existing SSSIs, already protected for other purposes, is certainly appropriate if their supporting habitat occurs within their boundaries. This is the case for the populations in lochs Maree and Calder, for example. Further, protection of supporting habitats through Water Framework Directive (WFD) River Basin Management Plans, which focus on water regulation and planning activities, and the application of clear regulation to prevent the introduction of non-native species can all contribute to the health and security of existing populations (SEPA, 2021).

Our results strongly urge against the movement of Arctic charr between lakes in different river catchments particularly those on the opposite sides of the east-west drainage divide, in alignment with previous suggestions for the conservation of the species (Maitland et al., 2007), due to the lack of contemporary connection between populations. In other Scottish salmonids of conservation concern, such as European whitefish (powan), translocations have been used with good success (Adams et al., 2014; Crotti et al., 2021; Lyle et al., 2006). If translocations are applied in Arctic charr, then we suggest that population genetic patterns should guide the practice with regard to population profiles. At southern portions of the species range in the UK, where populations are of high concern due to rapid declines, translocations are considered as a possible feature of conservation strategy and have already been applied to Loch Doon (Lavictoire et al., 2025; Maitland et al., 2007). The general high levels of genetic differentiation suggest that conservation of most populations need to be handled independent of one another.

## Conclusions

Populations clustered into their river catchments and on a wider scale by river system flow however levels of genetic differentiation were high, even between populations in the same river systems, suggesting low levels or a lack of gene flow and that most populations are reproductively isolated from one another. Identification of putatively adaptive SNPs highlighted a number of genes and regions with roles in local adaptation that warrant further investigation to determine the importance of their role. Our metrics of population vulnerability highlighted the value of using multiple metrics to explore vulnerability demonstrating who the geographical location and size of lake environment can affect population vulnerability. We focused conservation efforts on populations showing important phenotypes to preserve, such as ecological versatility, as well as that show important adaptive potential. Our study highlights the importance of wide-scale datasets when delineating conservation units as well as the challenging for appropriate delineations in highly diverse species.

## Acknowledgements

The authors thank Eric Verspoor and Ron Greer for their help in collecting and providing samples for 12 of the Scottish lochs included in this dataset; and Peter Koene, Hannele Honkanen, Rowan Smith, Jamie Carruth, Sandy Macfarlane and Ruaidhri Forrester for assistance with fieldwork. We thank David Wood and the RSPB Oa reserve team with facilitating access to Loch Ard Achadh, and Alastair Stephen for support with the applied project development. Thanks to Maria Capstick for teaching the lab protocols used in this project and Kara Layton for her help and insight with the genetic offset analysis. Thanks also to Arne Jacobs for providing existing data for eight of the studied lakes and with the bioinformatic processing and analysis. This research was supported by NatureScot, SSE, and the School of Biodiversity, One Health & Veterinary Medicine at the University of Glasgow.

## Data accessibility and benefits-sharing

Data accessibility: Relevant sample meta-data, environmental information, SNP VCF files, and code used in the analyses are archived on the University of Glasgow’s permanent data repository, Enlighten (doi with acceptance). Demultiplexed ddRADseq data are available on NCBI Short Read Archive (SRA) (BioProjects PRJNA1061680, PRJNA607173).

Benefits Generated: A research collaboration was developed with scientists and fisheries/conservation managers and stakeholders. All collaborators who provided material and feedback and/or collections support are included as co-authors or in acknowledgements. The results of research are shared publicly, with the relevant conservation communities, and the broader scientific community. This research addresses a priority concern for Scotland, in this case the conservation of organisms being studied. Benefits from this research accrue from the sharing of our data and results on public databases as described above.”

