## Supplementary Document for "Defining conservation units in a highly diverse species: A case on Arctic charr"

Defining conservation units in a highly diverse species: A case study identifying priority populations of Arctic charr

**Table of Contents:**

| **Figure S1** | Page 2 |
| --- | --- |
| **Figure S2** | Page 3 |
| **Figure S3** | Page 4 |
| **Figure S4** | Page 5 |
| **Figure S5** | Page 6 |
| **Figure S6** | Page 7 |
| **Figure S7** | Page 8 |
| **Figure S8** | Page 9 |
| **Figure S9** | Page 10 |
| **Table S2** | Page 11 |
| **Table S4** | Page 12 |
| **Table S6** | Page 12 |
| **Table S7** | Page 13 |
| **Table S9** | Page 13 |
| **Table S12** | Pages 14-15 |
| **Table S13** | Page 15 |

Figure S1.

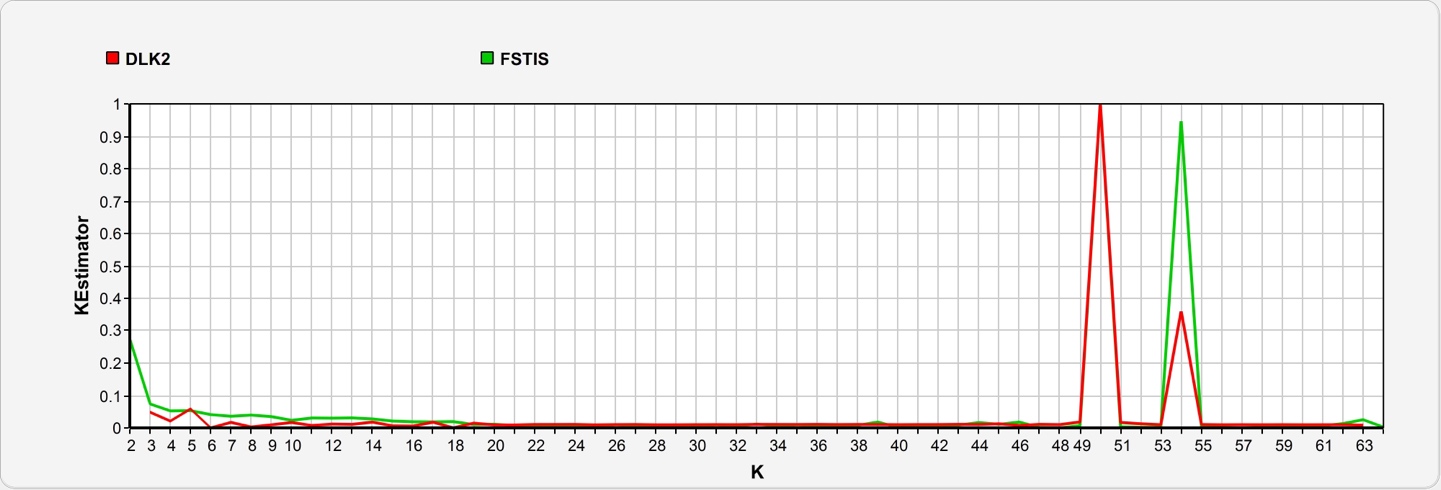

Figure S1. Plots showing the values of two estimators DLK2 and FSTIS estimates across the various K (number of genetic clusters) tested in admixture analysis to determine the most likely number of K. Higher values for each indicator suggest a more probable K value. Detailed explanation of how the estimators are calculated can be found in Wang et al. 2022.

Figure S2.

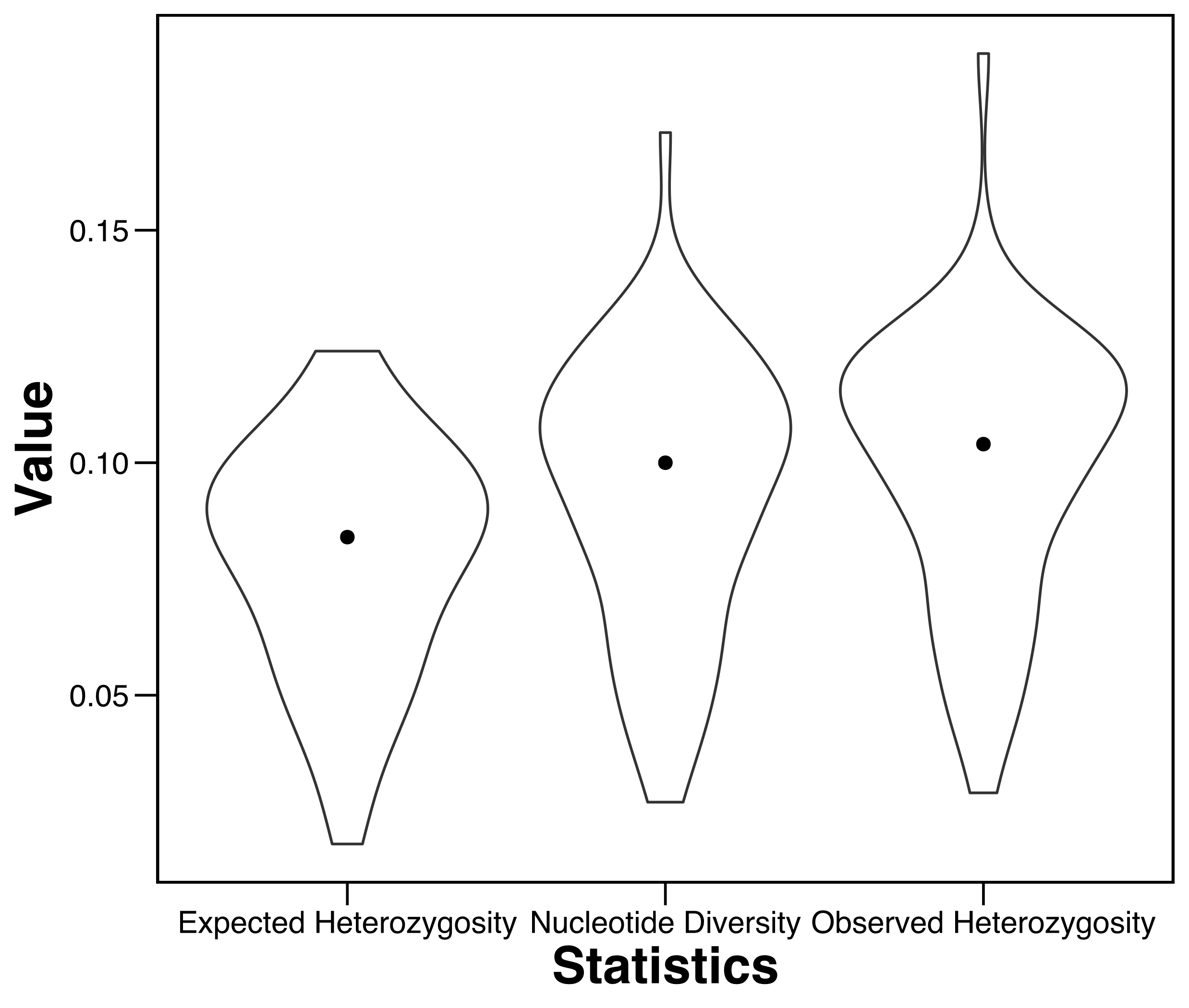

Figure S2. Violin plot of Expected Heterozygosity, Nucleotide Diversity and Observed Heterozygosity ranges for all populations. Black dot represents the mean values.

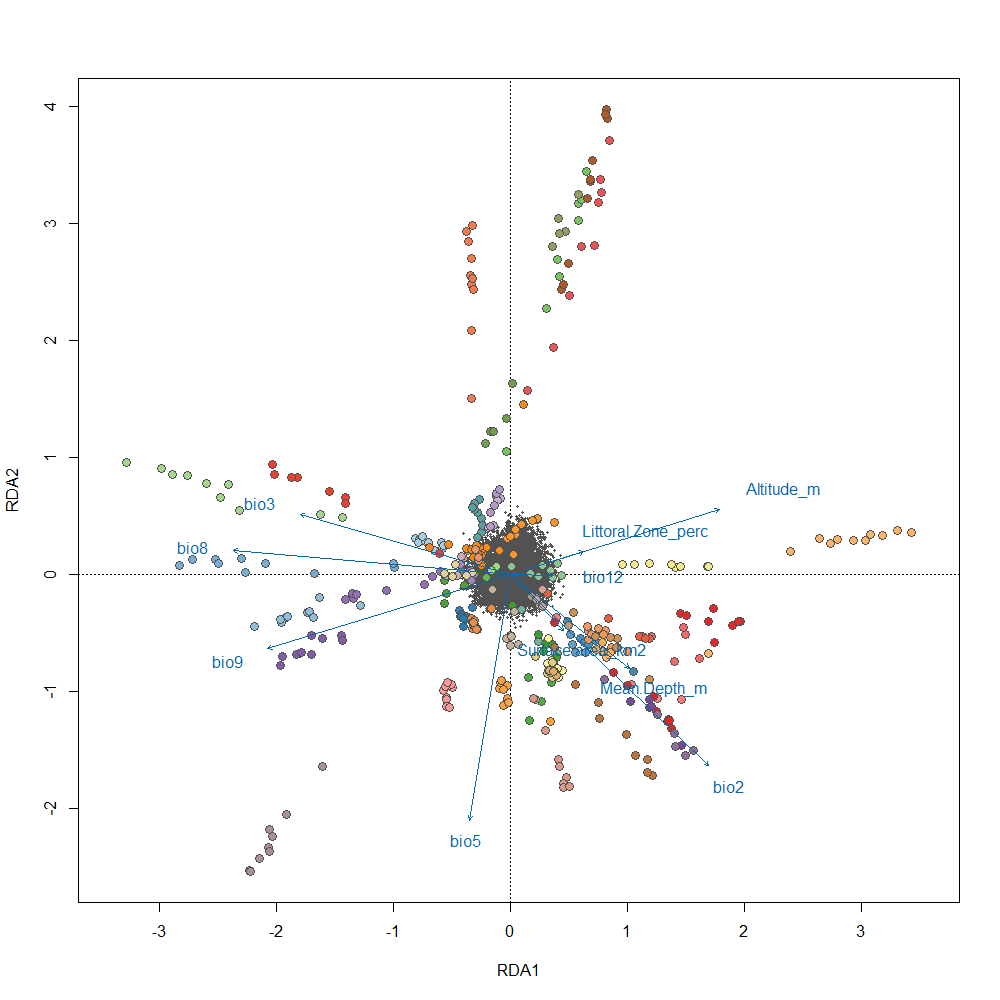
Figure S3.

Figure S3. Redundancy analysis plot showing the directions each uncorrelated variables pull in across RDA1 and RDA2. Colours represent each of the individual populations in the dataset.

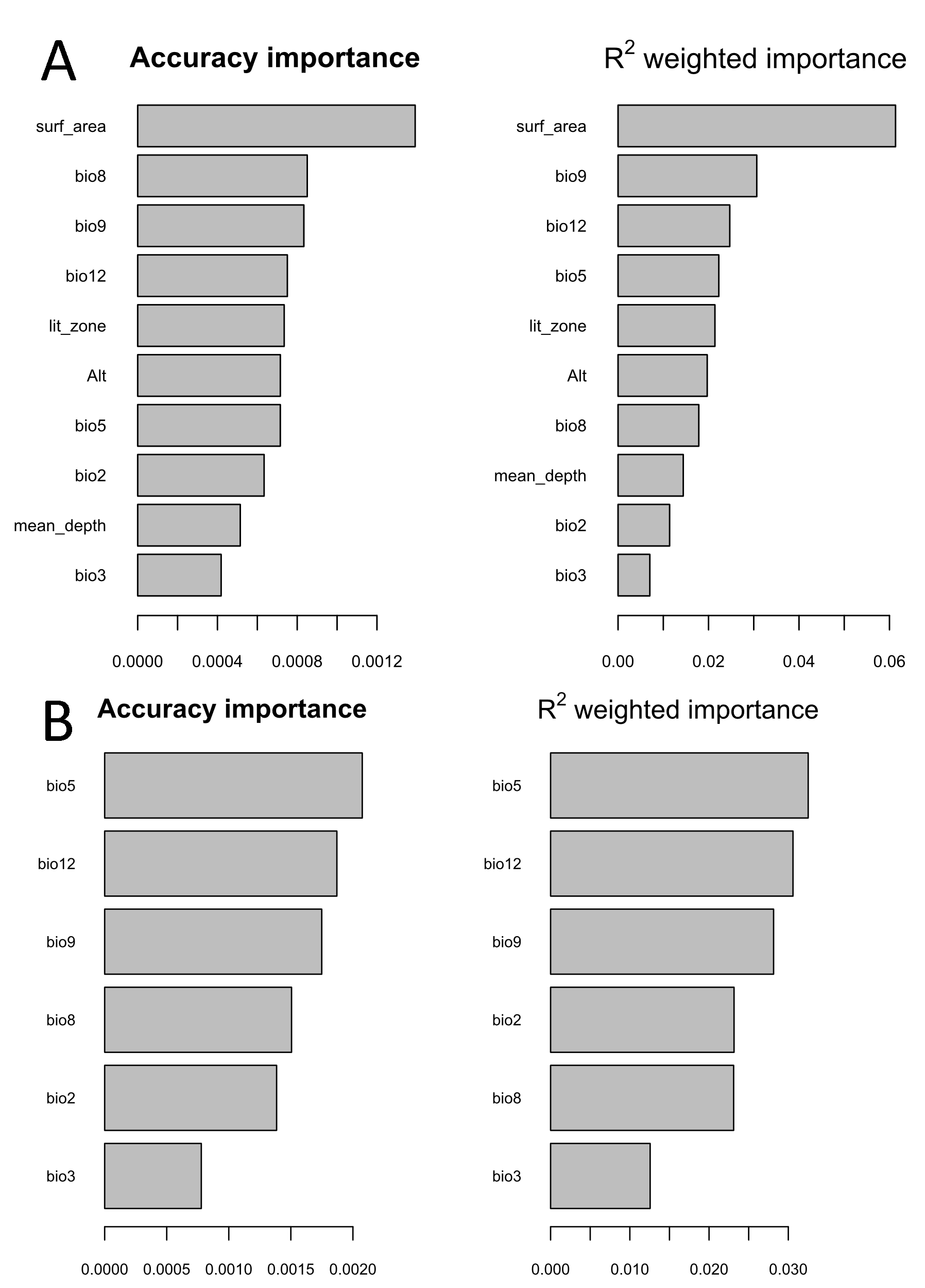
Figure S4.

Figure S4. R-squared importance plots for uncorrelated environmental variables in explaining A) adaptive variance (N=1,104 SNPs) and B) variance of climate associated SNPs used for genetic offset analysis (N=235 SNPs).

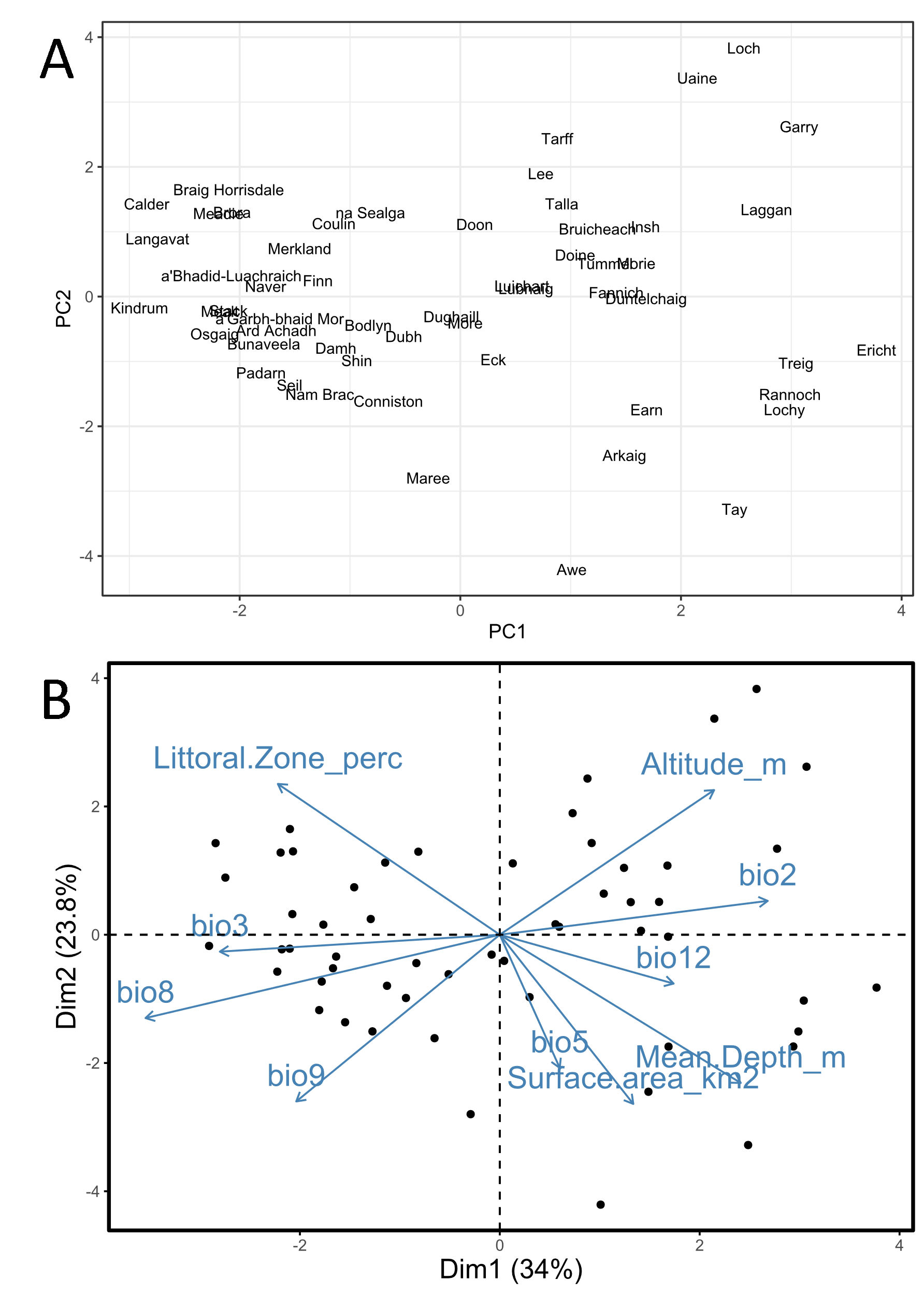
Figure S5.

Figure S5. Principal component analysis run on the 10 uncorrelated environmental variables used to identify the putatively adaptive SNPs. A) shows the placements of each lake of origin on PC1 and PC2 while B) shows the corresponding loadings of each of the environmental variables.

Figure S6.

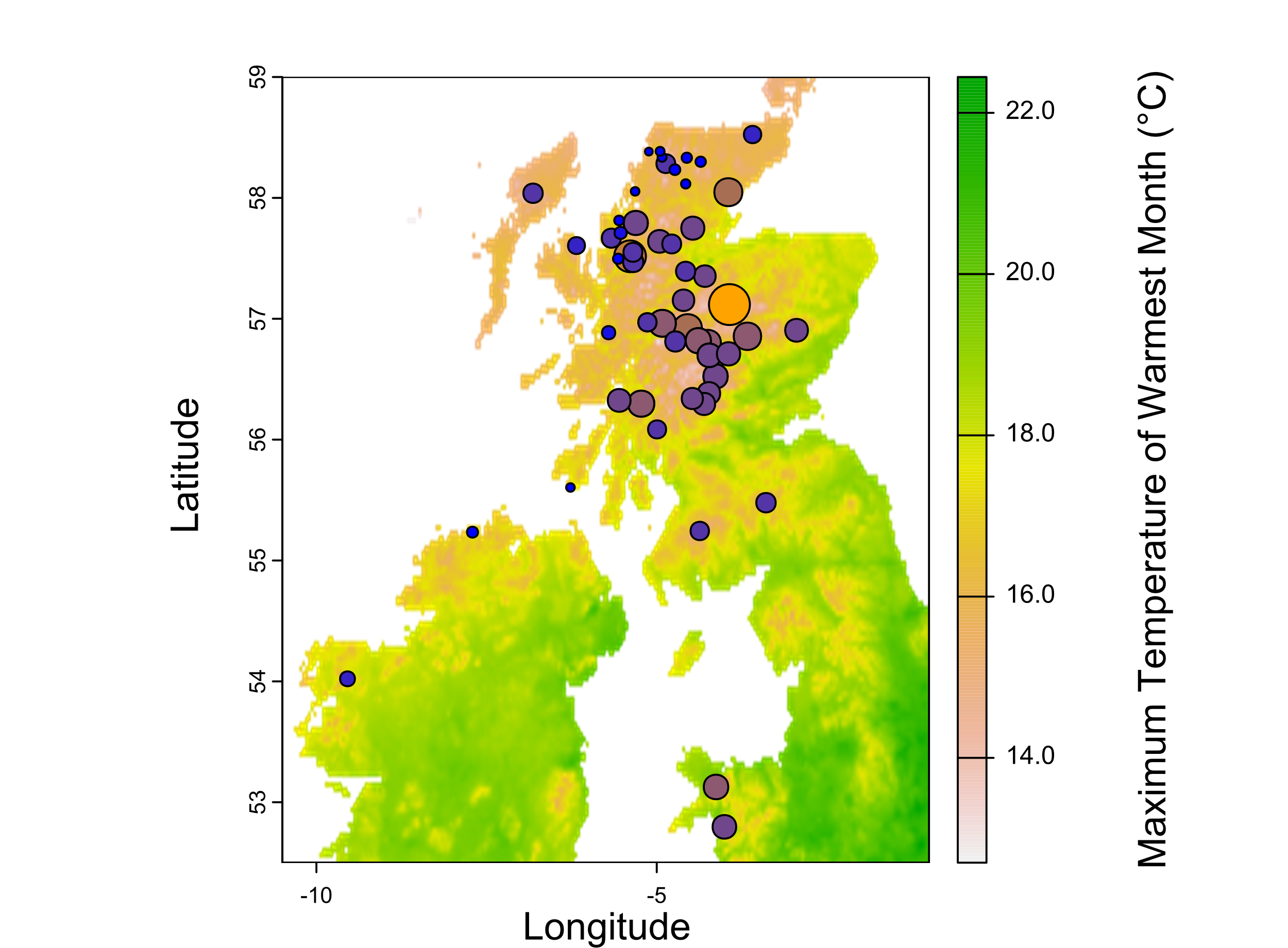

Figure S6. Plot of geographic distribution of populations. Circle sizes and colour reflect genetic offset scores (under the RCP 4.5 scenario) with larger and more orange ones representing a higher offset score. Map of the British Isles is coloured by Bio5 (Maximum temperature of the Warmest Month) readings.

Figure S7.

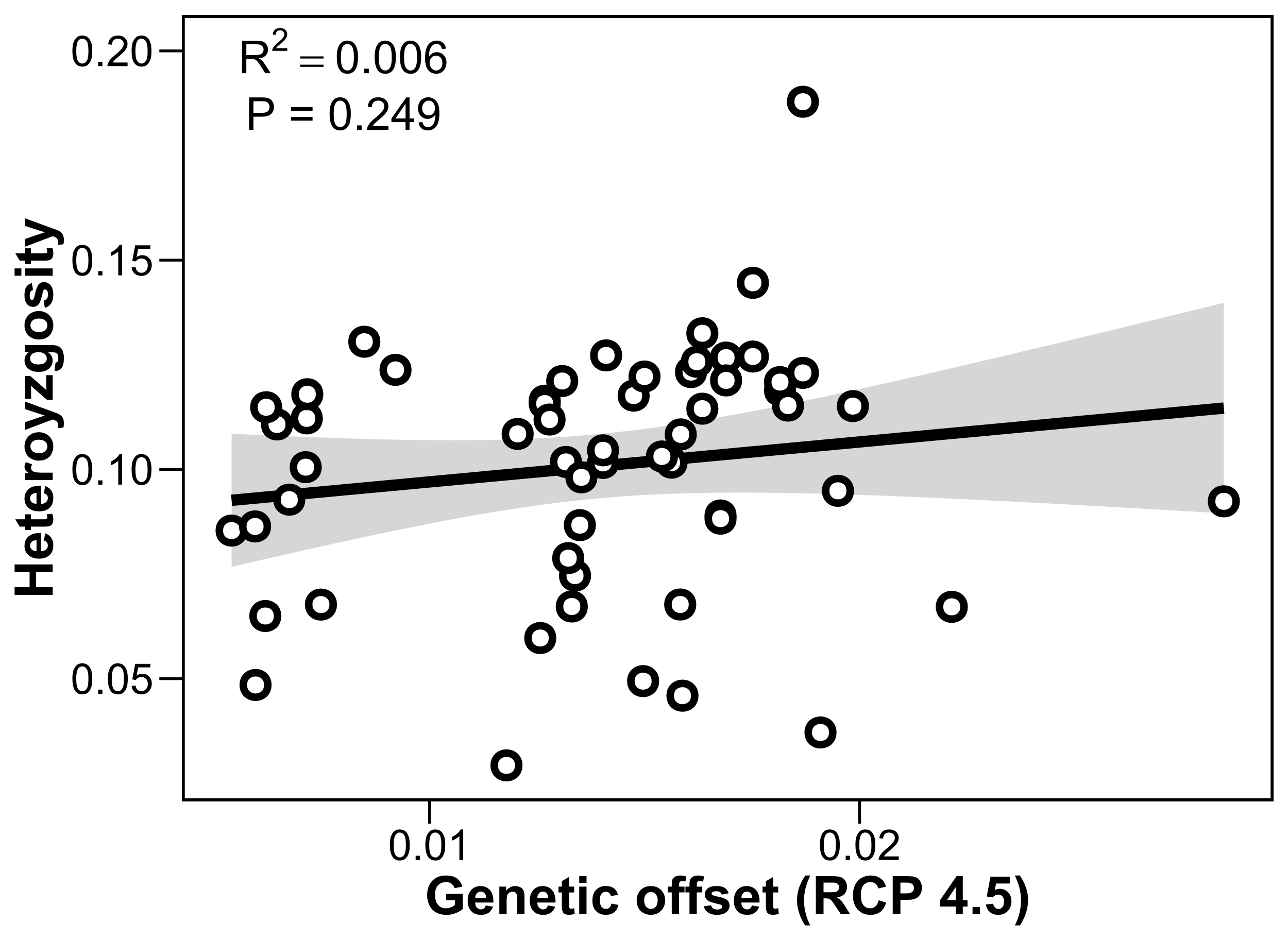

Figure S7. Relationship between genetic offset scores and levels of observed heterozygosity. Populations in the same lake of origin (i.e. ecotype pairs) are given the same offset score.

Figure S8.

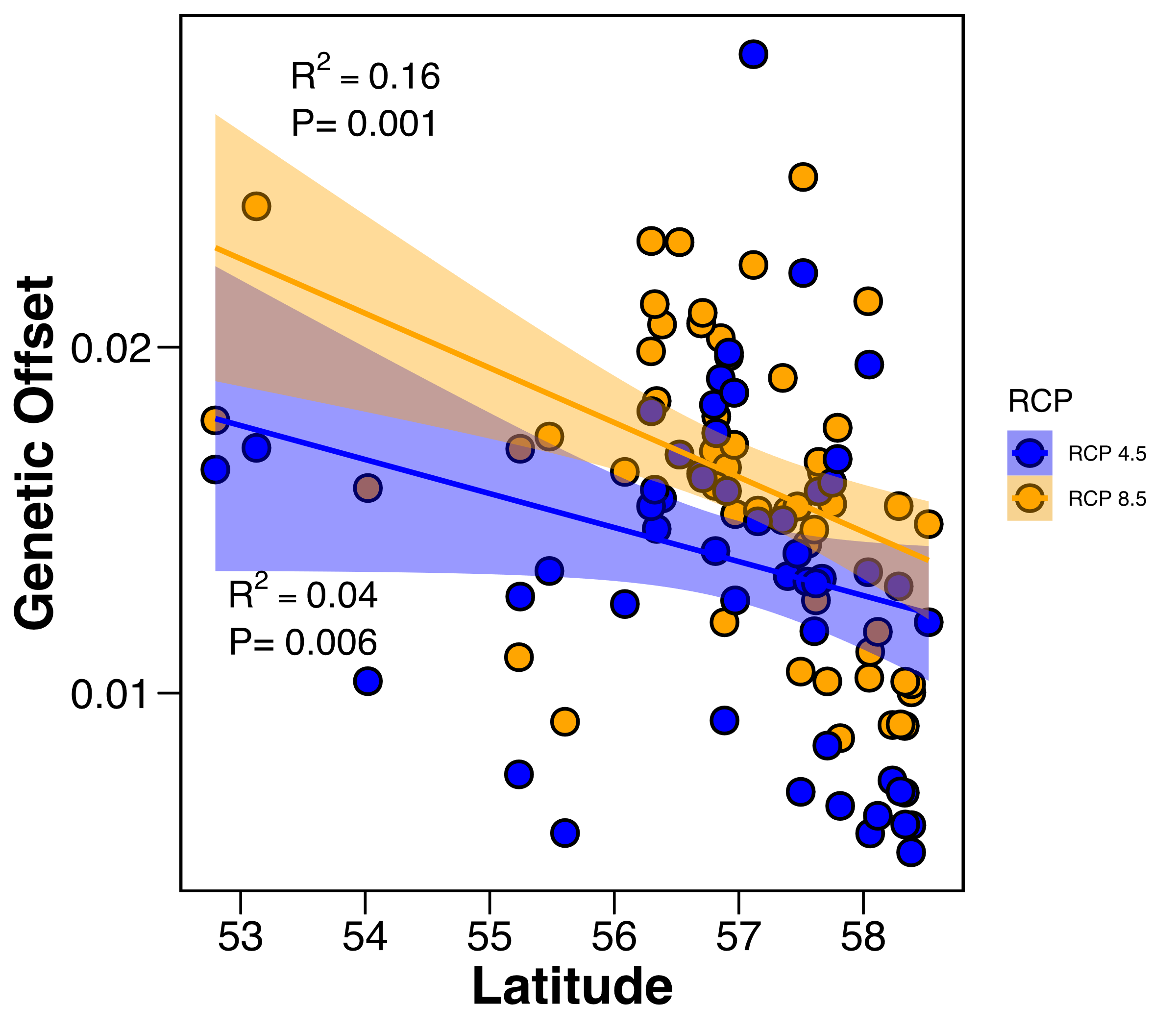

Figure S8. Genetic offset scores vs latitude under two RCP scenarios, RCP 4.5 (in blue) and RCP 8.5 (orange). Each point represents a different lake of origin.

Figure S9:

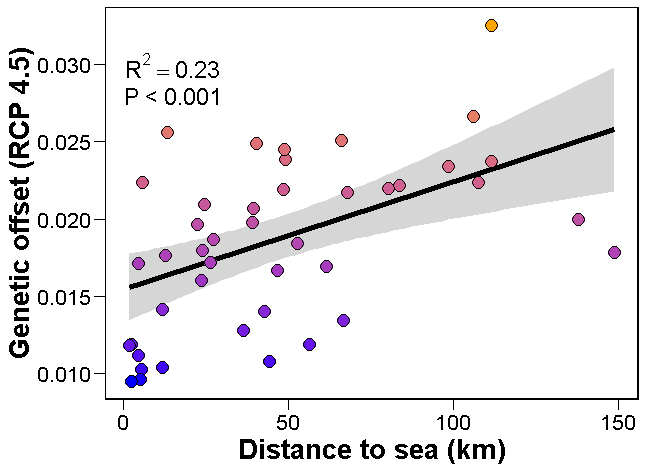

Figure S9: The relationship between genetic offset and distance to sea when only using populations with five or more individuals when calculating genetic offset.

Table S2.

Table S2. List of the 19 climatic variables and 5 bathymetric variables used in the study. Code and full names are provided. Variables in bold are the 10 uncorrelated variables used to identify adaptive SNPs.

| **Code** | **Variable** |
| --- | --- |
| Bio1 | Annual mean temperature |
| **Bio2** | **Mean diurnal range** |
| **Bio3** | **Isothermality (Bio2/Bio7)** |
| Bio4 | Temperature Seasonality |
| **Bio5** | **Maximum temperature of the warmest month** |
| Bio6 | Minimum temperature of the coldest month |
| Bio7 | Temperature annual range (Bio5-Bio6) |
| **Bio8** | **Mean temperature of wettest quarter** |
| **Bio9** | **Mean temperature of driest quarter** |
| Bio10 | Mean temperature of warmest quarter |
| Bio11 | Mean temperature of coldest quarter |
| **Bio12** | **Annual precipitation** |
| Bio13 | Precipitation of wettest month |
| Bio14 | Precipitation of driest month |
| Bio15 | Precipitation seaonality |
| Bio16 | Precipitation of wettest quarter |
| Bio17 | Precipitation of driest quarter |
| Bio18 | Precipitation of warmest quarter |
| Bio19 | Precipitation of coldest quarter |
| **Alt** | **Altitude** |
| Max.Depth | Maximum lake depth |
| **Mean.Depth** | **Mean lake depth** |
| **S.Area** | **Lake surface area** |
| **Lit.Zone** | **Percentage of the lake that is < 5 metres deep (Littoral zone)** |

Table S4.

Table S4. Scoring system used to calculate lake sensitivity scores based on lake surface area, depth, and altitude.

| **Max depth (m)** | **Depth sensitivity score** | **Surface Area (ha)** | **Surface Area sensitivity score** | **Altitude (m)** | **Altitude sensitivity score** |
| --- | --- | --- | --- | --- | --- |
| <10 | 4 |  |  | <10 | 4 |
| 10 to 30 | 3 | <10 | 3 | 10 to 50 | 3 |
| 30 to 100 | 2 | 10 to 30 | 2 | 50 to 200 | 2 |
| >100 | 1 | >30 | 1 | >200 | 1 |

Table S6.

Table S6. Hierarchical AMOVA results showing the variance explained by various groupings including the east-west drainage divide seen in the neighbour-joining tree and hydrometric area of origin.

|  | **Degrees of freedom** | **Sum squares** | **Mean squares** |
| --- | --- | --- | --- |
| Between East-West split | 1 | 7017.08 | 7017.08 |
| Between Hydrometric Areas within East-West split | 30 | 117095.98 | 3903.20 |
| Between Populations within Hydrometric Areas | 32 | 68307.97 | 2134.62 |
| Between Samples within Populations | 345 | 69268.06 | 200.20 |
| Within Samples | 410 | 108513.12 | 264.67 |
| Total | 819 | 370202.21 | 451.02 |

|  | **Sigma** | **%** |
| --- | --- | --- |
| Variation between East-West split | 2.54 | 0.55 |
| Variation between Hydrometric Areas within East-West split | 62.39 | 13.56 |
| Variation between Populations within Hydrometric Areas | 162.68 | 35.36 |
| Variation between Samples within Populations | -32.23 | -7.01 |
| Variation within Samples | 264.67 | 57.53 |
| Total variation | 460.04 | 100.00 |

|  | **Phi** |
| --- | --- |
| Phi-Samples-total | 0.43 |
| Phi-Samples-Populations | -0.14 |
| Phi-Populations-Hydrometric Areas | 0.41 |
| Phi-Hydrometric Areas-East West split | 0.14 |
| Phi-East West split-total | 0.01 |

Table S7.

Table S7. ANOVA results showing the significance of each constrained axis in the RDA in explaining variance

|  | **DF** | **Variance** | **F** | **Pr (>F)** |  |
| --- | --- | --- | --- | --- | --- |
| RDA1 | 1 | 459.2 | 8.3956 | 0.001 | *** |
| RDA2 | 1 | 446.3 | 8.1611 | 0.001 | *** |
| RDA3 | 1 | 387.3 | 7.0814 | 0.001 | *** |
| RDA4 | 1 | 351.8 | 6.4328 | 0.001 | *** |
| RDA5 | 1 | 307.7 | 5.6268 | 0.001 | *** |
| RDA6 | 1 | 292.4 | 5.3457 | 0.001 | *** |
| RDA7 | 1 | 261.8 | 4.7869 | 0.001 | *** |
| RDA8 | 1 | 213.7 | 3.9076 | 0.001 | *** |
| RDA9 | 1 | 183.6 | 3.3562 | 0.001 | *** |
| RDA10 | 1 | 152.1 | 2.7815 | 0.001 | *** |
| Residual | 399 | 21822.1 |  |  |  |

Table S9.

Table S9. Number of putatively adaptive SNPs (N=1,104) each uncorrelated environmental variables was the strongest predictor for in the Redundancy Analysis.

| **Predictor** | **Number of SNPs** |
| --- | --- |
| Surface area | 234 |
| Mean depth | 70 |
| Littoral zone percentage | 35 |
| Altitude | 50 |
| Bio2 | 166 |
| Bio3 | 156 |
| Bio5 | 127 |
| Bio8 | 29 |
| Bio9 | 166 |
| Bio12 | 71 |

Table S12.

Table S12. List of QTLs from Fenton *et al.* 2024b that contained adaptive SNPs we identified. QTL marker names are taken from the mentioned paper. Species of origin and trait the QTL is associated with are also listed. Chromosome, QTL start, and end refer to QTL location when mapped to the *Salvelinus sp.* genome.

| **QTL marker** | **Species** | **Trait** | **Chromosome** | **QTl start** | **QTl end** |
| --- | --- | --- | --- | --- | --- |
| Cocl_BS_096 | *Coregonus clupeaformis* | Body Shape | NC_036838.1 | 46685809 | 46685809 |
| Omy_SM_031 | *Oncorhynchus mykiss* | Sexual maturation | NC_036838.1 | 50898901 | 50898901 |
| Cocl_GR_004 | *Coregonus clupeaformis* | Directional change | NC_036839.1 | 10561411 | 10561411 |
| Cocl_BS_021 | *Coregonus clupeaformis* | Body Shape | NC_036842.1 | 34066589 | 34066589 |
| Sal_CF_011 | *Salvelinus alpinus* | Condition Factor | NC_036842.1 | 5751355 | 88198010 |
| Cocl_DirCh_008 | *Coregonus clupeaformis* | Directional change | NC_036842.1 | 11649948 | 11649948 |
| Omy_DR_003 | *Oncorhynchus mykiss* | Disease Resistance | NC_036842.1 | 5127503 | 5127503 |
| Cocl_BS_053 | *Coregonus clupeaformis* | Body Shape | NC_036843.1 | 22695609 | 22695609 |
| Omy_SM_014 | *Oncorhynchus mykiss* | Sexual maturation | NC_036845.1 | 29414044 | 29414200 |
| Oki_BL_009 | *Oncorhynchus kisutch* | Body Length | NC_036847.1 | 24097769 | 32996863 |
| Omy_DR_012 | *Oncorhynchus mykiss* | Disease Resistance | NC_036848.1 | 13363254 | 13363254 |
| Sal_UTT_004 | *Salvelinus alpinus* | Upper temperature tolerance | NC_036848.1 | 10119484 | 21971402 |
| Omy_BW_016 | *Oncorhynchus mykiss* | Body Weight | NC_036853.1 | 10667932 | 17403275 |
| Omy_CF_009 | *Oncorhynchus mykiss* | Condition Factor | NC_036853.1 | 10667932 | 17403275 |
| Oki_SP_005 | *Oncorhynchus kisutch* | Spawning date | NC_036853.1 | 10667932 | 17403275 |
| Sna_BS_006 | *Salvelinus namaycush* | Body Length | NC_036854.1 | 3663455 | 3663455 |
| Sna_BL_133 | *Salvelinus namaycush* | Body Length | NC_036855.1 | 50063281 | 50063281 |
| Sna_BL_043 | *Salvelinus namaycush* | Body Length | NC_036855.1 | 32096909 | 32096909 |
| Omy_OSMO_005 | *Oncorhynchus mykiss* | Sodium/Chloride concentration 2 | NC_036855.1 | 17674273 | 17674527 |
| Omy_BW_031 | *Oncorhynchus mykiss* | Body Weight | NC_036856.1 | 18229605 | 18229605 |
| Omy_SP_018 | *Oncorhynchus mykiss* | Spawning date | NC_036856.1 | 42026512 | 42027014 |
| Sal_CF_095 | *Salvelinus alpinus* | Condition Factor | NC_036861.1 | 875380 | 875380 |
| Sal_HS_019 | *Salvelinus alpinus* | Head Shape | NC_036861.1 | 875380 | 875380 |
| Oki_SP_011 | *Oncorhynchus kisutch* | Spawning date | NC_036862.1 | 27196736 | 35780323 |
| Sna_BL_221 | *Salvelinus namaycush* | Body Length | NC_036863.1 | 42690182 | 42690182 |
| Sfo_GR_005 | *Salvelinus alpinus* | Growth Rate | NC_036865.1 | 12786571 | 12786571 |
| Sal_BW_072 | *Salvelinus alpinus* | Body Weight | NC_036866.1 | 14035337 | 44737836 |
| Cocl_DirCh_001 | *Coregonus clupeaformis* | Directional change | NC_036869.1 | 19718333 | 19718333 |
| Cocl_DirCh_005 | *Coregonus clupeaformis* | Directional change | NC_036870.1 | 5780669 | 5780669 |
| Oki_SM_002 | *Oncorhynchus kisutch* | Sexual maturation | NC_036871.1 | 12958985 | 12958985 |
| Sna_BS_058 | *Salvelinus namaycush* | Body Length | NW_019942665.1 | 366314 | 366314 |
| Cocl_BS_169 | *Coregonus clupeaformis* | Body Shape | NW_019942665.1 | 364528 | 364528 |
| Cocl_BS_167 | *Coregonus clupeaformis* | Body Shape | NW_019943451.1 | 91315 | 91315 |
| Cocl_BS_101 | *Coregonus clupeaformis* | Body Shape | NW_019943516.1 | 79438 | 79438 |
| Sal_BL_005 | *Salvelinus alpinus* | Body length | NW_019945303.1 | 61402 | 61582 |

Table S13.

Table S13. Correlation matrix between the different metrics of population vulnerability: genetic offset scores, lake sensitivity scores, and observed heterozygosity.

|  | **Genetic offset** | **Sensitivity score** | **Heterozygosity** |
| --- | --- | --- | --- |
| Genetic offset | 1.00 | -0.40 | 0.26 |
| Sensitivity score | -0.40 | 1.00 | -0.45 |
| Heterozygosity | 0.26 | -0.45 | 1.00 |
